# Transient species can increase resident metapopulation size by modifying local conditions and regional connectivity

**DOI:** 10.64898/2026.01.07.698263

**Authors:** William Jia-Ang Ou, Rachel M. Germain, Tadashi Fukami

## Abstract

Populations are regulated by processes operating at both local and regional scales. As a result, species can interact not only within local communities but also through regional-scale processes. However, the feedback between local and regional processes of species interactions in shaping metapopulation dynamics have rarely been tested experimentally. Using nectar microbes as a model system, we examine how a bacterial species dispersing to a sink habitat, sustained by external supply, can interact with a resident yeast species directly by modifying local nectar chemistry and indirectly by altering metapopulation connectivity. We show that the external supply of bacteria increases resident yeast densities, potentially through increased local productivity, and that this benefit is enhanced when the dispersal rate of yeast is reduced in patches supplied with bacteria. Analysis of a metapopulation model indicates that the benefit of dispersal modification on metapopulation size occurs only when local productivity differs across patches, consistent with our interpretation that bacterial supply increases local productivity. Taken together, our results underscore the role of dispersal in shaping how metapopulations experience density-dependence across heterogeneous landscapes, highlight the relevance of material flows across ecosystem boundaries, and suggest a plausible mechanism in which transient species may leave lasting impacts on ecological communities.

## INTRODUCTION

Many natural populations are spatially subdivided into a network of local populations connected by dispersal, *i.e.*, a metapopulation. Decades of theoretical work has shown that dispersal does more than merely shuffle individuals across space (Hastings, 1982; Holt, 1985; Karlin & McGregor, 1972; Levin, 1974; Levins, 1969). Even under the simplest scenarios, dispersal of organisms can have non-intuitive effects on local and regional patterns of biodiversity (Hanski, 1998; Huffaker, 1958; Levin, 1974; Levins, 1969). For instance, dispersal enables metapopulations to persist both through patch dynamics, when local populations fluctuate to extinction (Brown & Kodric-Brown, 1977), and through inflationary effects, when occasional bouts of positive growth offset negative average local growth rates (Kortessis et al., 2025; Matthews & Gonzalez, 2007). Recent work demonstrates that dispersal can even increase the total number of individuals across the metapopulation (i.e., metapopulation size) relative to the sum of its local carrying capacities without dispersal if local conditions are spatially heterogeneous (Arditi et al., 2015; DeAngelis et al., 2016; He et al., 2025; Zhang et al., 2017). In sum, these results indicate that the dynamics of a regional metapopulation are rarely the simple sum of its constituent populations’ dynamics considered in isolation, but instead depend on how dispersal links them.

Despite progress made in understanding how dispersal structures ecological communities, relatively less is known about the ecological consequences when the dynamics of dispersal is itself coupled to local dynamics. In natural ecological settings, non-random (directed or biased) dispersal is more likely the norm than the exception (Amarasekare, 2010; Clobert et al., 2009; McPeek et al., 2024; Vuilleumier & Possingham, 2006). While directionality in movement can arise naturally via physical forces such as wind, currents, and gravity (Germain et al., 2019; Mari et al., 2014; Okubo & Levin, 2001), it can also result from biotic mechanisms such as habitat selection (Clobert et al., 2009). Although it might seem that directed dispersal via habitat selection is unique to mobile animals with cognitive capacities, organisms with limited motility can also disperse non-randomly via mechanisms such as phoresy (or zoochory in plants), where their interactions with animal dispersal vectors can dictate both the rate and direction of movement (Bartlow & Agosta, 2021). Given that dispersal can modify population size and that species can influence one another’s dispersal, it follows that species interactions can be mediated through changes in dispersal. For example, phoretic species that share limited dispersal vectors could compete for them in ways that shape their relative abundances at the metapopulation level (Nell et al., 2025), yet few empirical studies have addressed this.

Recognising that dispersal can be an integral component of how species interact could reveal cryptic interactions that shape ecological communities. This possibility is particularly relevant for species that form sink populations—i.e., species whose persistence is determined less by local conditions than by immigration from adjacent habitats. Because such species cannot maintain growth locally, their potential impacts on ‘resident’ species—i.e., species capable of persisting without immigration—are often overlooked (Amor et al., 2020; Mallon et al., 2018). However, previous studies suggest that sink populations are not only prevalent but can also impact resident species (Amarasekare & Nisbet, 2001; Holt et al., 2003; Long et al., 2007; Mallon et al., 2018; Pulliam, 1988). This impact occurs primarily through the sheer abundance of individuals immigration introduces, amplifying their total impact even if their per-capita effects on residents is otherwise small. For instance, a species occupying a sink habitat can reach sufficiently high densities via immigration and deplete resources to levels that depress, or even competitively exclude, an otherwise locally superior competitor (Amarasekare & Nisbet, 2001; Long et al., 2007). Notably, these studies assume that sink populations impact residents purely through local interactions. However, if dispersal can be shaped by other species, sink populations may also influence residents by altering their dispersal patterns across the metapopulation. Despite evidence showing that shared dispersal vectors can reshape dispersal patterns among competing species (e.g., Muller-Landau et al., 2008; Xiao & Zhang, 2016), their long-term demographic consequences remain unknown.

A system in which dispersal vectors modulate species interactions is the floral microbiome, making it an excellent model system for examining feedbacks between local and regional processes. Floral nectar often contains microbial communities (Vannette, 2020) that are dispersed by pollinators (Belisle et al., 2012; Herrera et al., 2009; Vannette & Fukami, 2017), whose identity and foraging patterns shape the spatial distribution of nectar microbes (Belisle et al., 2012; Morris et al., 2020; Vannette et al., 2021; Vannette & Fukami, 2017). Because these microbes are often a subset of the communities carried by pollinators, nectar likely contains many transient invaders (Mallon et al., 2018). Importantly, the microbial activity within nectar can alter pollinator foraging behaviour by modifying nectar chemistry or producing volatiles that attract or repel certain pollinators (Chappell et al., 2022; Schaeffer et al., 2019; Vannette et al., 2013). Such local modifications may bias dispersal towards or away from certain flowers, effectively enabling microbes to participate in their own dispersal despite being unable to move on their own. Together, these processes create a feedback loop between local dynamics and regional dispersal via pollinator foraging (Nell et al., 2025), providing an additional pathway in which sink populations could interact with residents at the metapopulation level.

Here, using nectar microbes isolated from the hummingbird-pollinated sticky monkeyflower (*Diplacus aurantiacus*), we conducted a laboratory experiment that simulates pollinator-assisted dispersal to investigate how a bacterial species invading a sink habitat can impact the metapopulation dynamics of a resident yeast species through local and regional processes. To avoid ambiguity, we refer to dispersal from external sources as “supply” (or “invasion”) and reserve “dispersal” for movement among patches within the metapopulation (external vs. internal dispersal *sensu* Fukami 2005). Previous work in this system has shown that the bacterial species reduces nectar pH, which in turn suppresses yeast growth and reduces pollinators’ nectar consumption (Chappell et al., 2022). In a pilot experiment, we found that the bacterial species cannot sustain a viable population under our experimental conditions but can nevertheless reduce nectar pH, making it well suited for examining the effects of species invading a sink habitat on resident yeast species. Specifically, we ask how the yeast metapopulation responds to changes in local nectar chemistry (i.e., reduced pH) and patch (*i.e*., local population) connectivity mediated by pollinator foraging preferences (simulated experimentally via pipetting) in the presence of bacterial sink populations. First, we test the hypothesis that the external supply of bacteria reduces yeast metapopulation size by lowering nectar pH to the detriment of local yeast growth (*i.e*., a local effect). Second, we test the hypothesis that pollinator preference biases dispersal away from low-pH patches, altering yeast metapopulation size by reshaping metapopulation connectivity (*i.e*., a local-regional interaction effect). As we show below, our findings ran opposite to these predictions. Bacterial supply increased yeast metapopulation size, and dispersal bias amplified this benefit. Because this reversal was unexpected, we analysed a simple two-patch model as a heuristic rather than a quantitative description of our system, to establish the conditions under which dispersal bias would amplify the local effects of an invading species on a resident’s metapopulation size.

## METHODS

### Overview of study system

To investigate the impact of a species invading a sink habitat on the metapopulation dynamics of a resident species, we conducted experiments using two nectar microbes: the bacterium *Acinetobacter nectaris* (strain FNA17) and the yeast *Metschnikowia reukaufii* (strain MR1). These microbes naturally inhabit the floral nectar of the sticky monkeyflower, *Diplacus aurantiacus,* and the specific strains we used were isolated from flowers in Jasper Ridge Biological Preserve (’Ootchamin ‘Ooyakma) located in the Santa Cruz Mountains of California. *Diplacus aurantiacus* is a shrub native to coastal Oregon and California and is commonly pollinated by hummingbirds (Belisle et al., 2012). Instead of dispersing between flowers themselves, these microbes “hitchhike” onto pollinators to move between flowers. When microbes arrive at a new flower, they proliferate by metabolising nectar resources and in so doing, modify nectar chemistry and set in motion the diverse ecological interactions that occur in these microscopic ecosystems (e.g., microbe-microbe interactions, plant-pollinator mutualisms; Chappell & Fukami, 2018; Quevedo-Caraballo et al., 2025). Its relative simplicity and tractability allows this study system to be replicated in laboratory settings with artificial nectar in microwell plates as flowers and pipettes to simulate pollinator-assisted dispersal (Chappell et al., 2022; Grainger et al., 2019; Letten et al., 2018; Tucker & Fukami, 2014; Vannette & Fukami, 2014). Following these prior studies, we adapted this experimental microcosm for the purpose of our study, detailed below.

### Experimental setup

We conducted an experiment that factorially manipulated ‘bacterial supply’ (i.e., the number of habitat patches in each metapopulation that would receive bacteria from an external source), dispersal bias, and dispersal frequency (i.e., the number of dispersal events within a growth cycle) to test how *A. nectaris* bacteria-driven pH changes affect *M. reukaufii* yeast metapopulation by modifying local nectar conditions and metapopulation connectivity through dispersal bias (Fig. 1). Note that our hypotheses are concerned with how patches are wired (*i.e.*, dispersal bias) and not the overall frequency of dispersal events *per se.* As such, we treated frequency as a scaling factor, rather than a focal variable, to ensure meaningful effect sizes of main factors could be detected. Because it was nonetheless a manipulated factor, we retained frequency and its two-way interactions in all models, both to account for the manipulated frequency treatment and to test, rather than assume, that the focal effects hold across dispersal rates.

**Figure 1.**
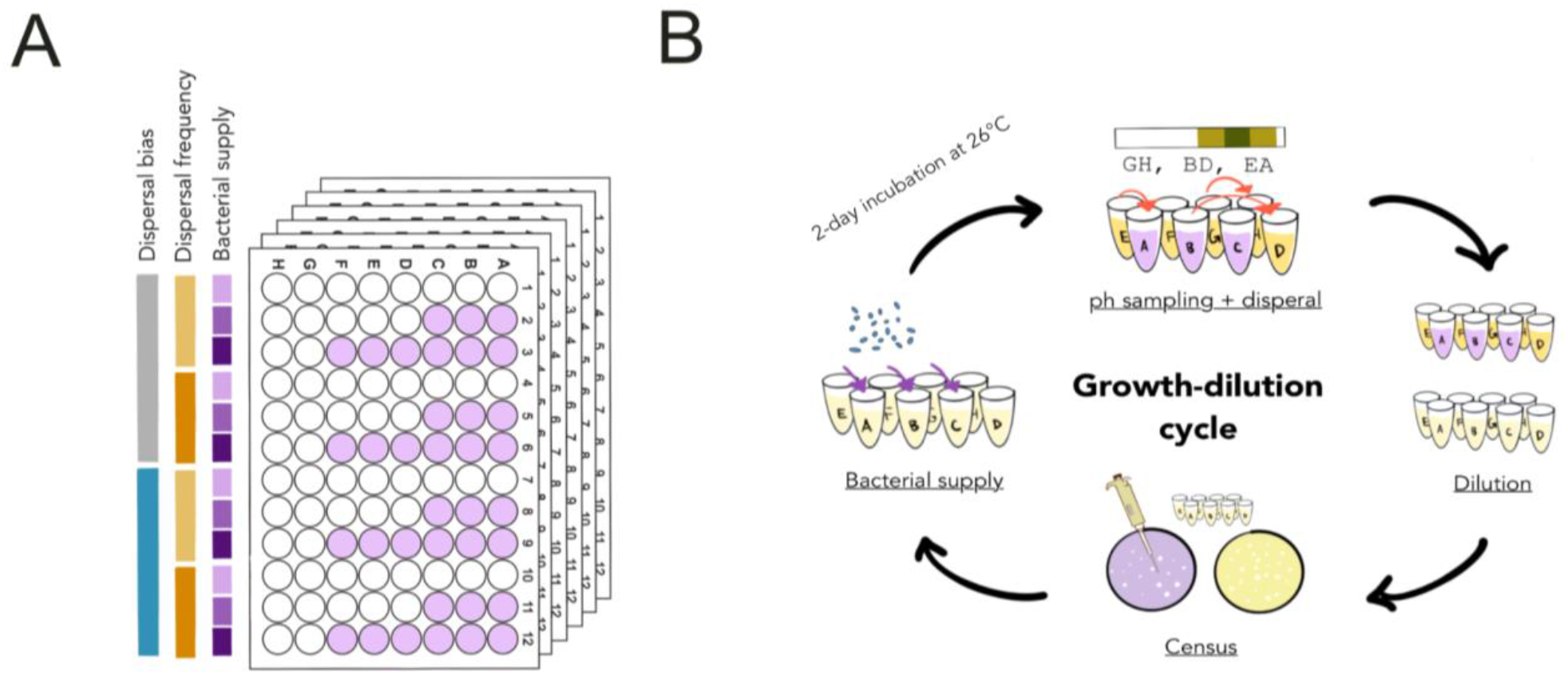
Overview of experimental setup. (**A**) Fully crossed design: dispersal bias (2) × dispersal frequency (2) × bacterial supply (3), each with six replicates. Each metapopulation consisted of 8 patches (PCR plate wells) connected by dispersal and underwent four 2-day growth–dilution cycles. (**B**) At the start of each cycle, patch pH was measured and used to generate donor–recipient pairs (see Methods). Dispersal was performed by transferring 3 µl of nectar from donor to recipient (red arrows). Patches were then diluted (5 µL into 95 µL fresh nectar) to emulate resource renewal, and samples were plated (TSA for bacteria, YMA for yeast) to estimate densities (CFUs µL^-1^). Bacteria (∼20,000 cells) were added according to treatment, and plates were sealed and incubated at 26°C for 2 days.

Our experimental metapopulations consisted of eight wells (i.e., patches) on a 96-well PCR plate, connected by dispersal. Each metapopulation was assigned to one of 12 treatment groups defined by the fully factorial combination (Fig. 1A) of bacterial supply (three levels: 0, 3, or 6), dispersal bias (two levels: random or bias), and dispersal frequency (two levels: 3 or 6), with each treatment replicated six times (12 treatments x 6 replicates = 72 metapopulations). Each replicate metapopulation was initiated by seeding patches with approximately 100-150 viable cells of *M. reukaufii* (see Supplementary Methods) and 100 µL of sterile artificial nectar. Artificial nectar was prepared as specified by Chappell et al. (2022) and the full recipe is provided in Supplementary Methods. Plates were sealed with a sterlised air-permeable membrane and incubated at 26°C for two days before beginning the experiment. After initiation, each metapopulation underwent a total of four two-day growth-dilution-cycles where each cycle included the following order of events: 1) pH sampling + dispersal, 2) resource replenishment, 3) community census, 4) bacterial supply, and 5) a two-day incubation period (Fig. 1B):

#### Event 1: Dispersal

Each growth-dilution cycle included two forms of dispersal, pollinator and background dispersal, both carried out by pipetting 3 µL of nectar from a donor patch to a recipient patch. The bias and frequency treatments simulated pollinator-assisted dispersal, whereas background dispersal provided a fixed baseline of connectivity common to every replicate metapopulation (e.g., simulating movement by wind or rain).

For pollinator-assisted dispersal, we generated a sequence of donor-recipient pairs and transferred nectar between them in sequential order. In the bias treatment, we first sampled patch pH by pipetting 10 µL of nectar from each patch onto pH indicator strips, then used the recorded values as pollinator-preference weights. Donor-recipient pairs were then drawn by sampling two patches from the eight (A–H) without replacement, with the probability of each patch being drawn proportional to its weight (using the sample() function in R). Specifically, the weight of patch *i* is *ω_i_* = *pH_i_*/(3 + *pH_i_*) with selection probability obtained by normalising 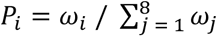 The half-saturation constant was set at three, which is the pH value that lies between nectar conditioned by bacteria (pH = 2) and yeast (pH = 4; Chappell et al. 2022). We mapped preference to pH via the Monod equation rather than assigning a fixed preference by patch type (*i.e.*, whether or not bacteria were present) because we did not know *a priori* how pH would be distributed across patches once dispersal occurred, nor how strongly bacteria would acidify the patches they occupied. This setup reflects the field observation that low-pH patches are less likely to be visited by pollinators and, consequently, microbes are less likely to disperse into or out of them (Vannette et al. 2013; Chappell et al. 2022). For the random (unbiased) treatment we skipped the pH-measurement step while still discarding 10 µL of nectar per patch, and drew donor-recipient pairs with equal probability across patches. The number of pollinator dispersal events per cycle was set by the frequency treatment (3 or 6), crossed factorially with dispersal bias.

For background dispersal, every replicate metapopulation received three dispersal events per cycle, drawn with equal probability across patches from a single pseudorandom sequence identical for all replicates. This ensured that patches within a metapopulation had a baseline level of connectivity that did not depend solely on dispersal by pollinators. Each replicate therefore underwent its 3 or 6 pollinator dispersal events plus 3 background events per cycle.

#### Event 2: Resource replenishment

Once the dispersal events were completed, all patches were diluted by transferring 5 µL of nectar into 95 µL of fresh, sterlised media on new 96-well plates to simulate resource replenishment (*i.e.*, nectar production in flowers). We used new sterile 96-well plates, instead of removing and adding nectar from the original, to minimise contamination.

#### Event 3: Sampling

Immediately after dilution, we sampled each local community by plating 5 µL of the newly diluted nectar onto yeast malt agar (YMA) for yeast and 5 µL onto tryptic soy agar (TSA) for bacteria, and then enumerated the resulting colony-forming units (CFUs) after >3 days of incubation at room temperature as a measure of microbial density (following Peay et al., 2012 and Vannette et al., 2013). Local densities or “population size” (CFU µL^-1^) were then calculated as the number of CFUs per sample volume (5 µL), while “metapopulation size” (CFU µL^-1^) was the average of local densities. Note that because local density and metapopulation size are both in units of CFU per unit volume, metapopulation size is the average and not the sum of local densities. Including the final census after the last cycle, each metapopulation was sampled five times. Six samples were lost to experimental error, resulting in 354 metapopulation-level observations.

Due to time constraints associated with simultaneously conducting the experiment and counting CFUs, we took images of the agar plates and enumerated CFUs after completion of the experiment (see Supplementary material for example images). In 35 of 354 metapopulation samples (9.89%), at least one local population was too numerous to count. Rather than discard these metapopulation samples, we assigned a value of 200 to the affected local populations, a value just above the maximum observed among countable local populations, before calculating metapopulation size. To ensure this artificial cutoff did not qualitatively affect our results, we repeated all analyses after excluding the 35 affected metapopulation samples (see ‘Potential artefacts of experimental design’ below).

#### Event 4: Bacterial supply

Following sampling, bacterial cells were added to patches according to the bacterial supply treatment. Specifically, bacterial cells were either added to no patches (bacterial supply = 0), patches A to C (bacterial supply = 3), or patches A to F (bacterial supply = 6). We chose to add bacteria to only a subset of patches within the metapopulation because this allows spatial heterogeneities in nectar pH and thus creates variation in dispersal probabilities under the dispersal bias treatment. Although doing so is not required, we chose to fix the patches in which bacteria were added (as opposed to randomly selecting patches to inoculate) to minimise experimental error. Approximately 20,000 bacterial cells (∼200 cells/µL nectar) were inoculated to each recipient patch (see Supplementary Methods). Bacterial cells were supplied after sampling because in our pilot trials, this amount of bacteria was necessary to reduce pH (after 2 days) but would be too dense to count if plated directly after inoculation. Thus, the bacteria that we sampled came from the remaining bacteria inoculated two days prior. Because bacterial inocula were prepared as a single batch for each growth cycle and distributed among all recipient patches, inoculum size was expected to vary more among cycles than among treatments within a cycle (Fig. S2). We account for this source of variation in the statistical analyses described below.

#### Event 5: Incubation

Following bacterial supply, 96-well plates were sealed with sterlised, air-permeable membranes and incubated at 26°C for 48 hours before beginning the next cycle

### Data analysis

#### Yeast metapopulation size

To assess treatment effects on yeast metapopulation size, we defined metapopulation size as the average of local patch densities (i.e., metapopulation density; CFU µL^-1^) rounded to the nearest integer and modelled it as a response variable in a negative binomial generalised linear mixed model (GLMM). Six samples (where ‘sample’ refers to a given metapopulation on a given sampling day) were excluded because pipetting errors left one or more patches unmeasured, giving *n* = 354 complete samples for analysis (Table S5). This model included dispersal frequency and bacterial supply as continuous predictors, dispersal bias as a categorical variable, and all two-way interactions as fixed effects, with replicate and sampling day as random effects in both the conditional and dispersion components. We included all two-way interactions because they aligned with our hypotheses and excluded the three-way interaction term because including it did not improve model fit. Because initial statistical fits with Poisson-distributed errors showed violation of homoscedasticity, we implemented a forward model selection to account for potential variance structure in our data. Specifically, we sequentially added treatment terms as predictors to the dispersion component of the negative binomial GLMM and only kept terms that significantly improved model fit (as evaluated by *χ*^2^ tests; Table S1). This led to the selection of a model that included bacterial supply as a predictor of the variance in yeast metapopulation size (Table 1; *ΔAIC* = -1.92, *P* = 0.048) and satisfied the assumption of residual homoscedasticity.

**Table 1.** GLMM results for yeast metapopulation size (n = 354). Estimates are marginal effect sizes (log scale) for the conditional and dispersion components, with 95% confidence intervals and standard errors. Term significance is from Type II Wald *χ*^2^ tests. Bold indicates *P* < 0.05. The Bias estimate is the biased-minus-random contrast. The interaction estimates are differences in marginal slopes: Bacterial supply:Bias is the difference in the supply slope between biased and random dispersal; Bacterial supply:Frequency is the change in the supply slope from frequency 3 to 6; Bias:Frequency is the difference in the frequency slope between dispersal treatments.

| Component | Term | Estimate (95% CI) | SE | $\chi^2$ (df) | P |
| --- | --- | --- | --- | --- | --- |
| Conditional | Bacterial supply | **0.028 (0.005, 0.052)** | **0.012** | **5.68 (1)** | **0.017** |
|  | Bias | 0.068 (-0.045, 0.180) | 0.057 | 2.02 (1) | 0.156 |
|  | Frequency | 0.011 (-0.026, 0.049) | 0.019 | 0.67 (1) | 0.414 |
|  | Bacterial supply:Bias | **0.056 (0.010, 0.102)** | **0.024** | **5.74 (1)** | **0.017** |
|  | Bacterial supply:Frequency | 0.045 (0.000, 0.091) | 0.023 | 3.76 (1) | 0.053 |
|  | Bias:Frequency | -0.040 (-0.114, 0.034) | 0.038 | 1.11 (1) | 0.291 |
| Dispersion | Bacterial supply | **-0.172 (-0.344, -0.001)** | **0.087** | **3.88 (1)** | **0.049** |

To ensure that our statistical results are robust to modeling decisions, we conducted several supplementary analyses. First, because bacterial supply had three levels (0, 3, and 6), we refitted the model treating supply as a categorical predictor to allow for a nonlinear response. The results were qualitatively unchanged, although statistical power was reduced because the categorical model requires estimating an additional parameter (Table S3). Moreover, the more parsimonious model with supply coded as a continuous predictor received stronger statistical support (*ΔAIC* = -6.16). Second, we tested whether treatment effects varied across the five growth cycles by fitting a model in which each treatment term interacted with sampling day. Model comparison indicated that the time-dependent treatment effect model was not supported (*χ*^2^ = 4.864, df = 6, *P* = 0.561), so we retained the original time-independent treatment effect model. Furthermore, we also found no residual temporal autocorrelation in the fitted time-independent model (Durbin-Watson *P* = 0.31), indicating that an autoregressive model was unnecessary. As a final robustness check, we split the data by sampling day and fitted the same model to each subset. Because bacterial inocula were prepared independently for each growth cycle, sampling day represented a source of between-cycle variation independent of the experimental treatments (Fig. S2). Consequently, statistical support for the effects of bacterial supply (and its interaction with dispersal bias) was less consistent across sampling days, compounded by the reduced sample size due to subsetting (Table S4). These supplementary analyses support our baseline model, which includes sampling day as a random effect to account for between-day variation while estimating treatment effects across the full time series.

We also examined the correlation between local yeast and bacterial densities as indirect evidence for a local effect of bacteria on yeast. Because variation in local bacterial density was the result of experimental noise rather than a randomised treatment, any effect of bacteria could only be inferred indirectly through association. To account for statistical non-independence arising from repeated measurements of patches, we used a stratified permutation test. Specifically, each block was a single metapopulation replicate on a single sampling day, and the test statistic was the Pearson correlation between yeast and bacterial density across patches within each block, averaged across blocks (thus preserving the replicate and sampling day groupings used as random effects in the GLMM). We then generated the null distribution by permuting yeast densities relative to bacterial densities within each block (*n_perm_* = 2000) and computed a one-sided *P*-value as the proportion of permuted test statistics greater than or equal to the observed correlation.

All statistical analyses were conducted using the ‘glmmTMB,’ ‘emmeans,’ and ‘DHARMa’ packages in R.

### Potential artefacts of experimental design

We conducted additional tests to confirm that our experimental results were not contingent on the choices of our experimental design. We describe three specific tests here in brief, and provide more detailed descriptions in the Supplementary Methods, with results shown in Figs. S3-S5. First, to ensure that ceiling our uncountable local populations at 200 did not artificially bias our results, we refitted the same statistical models with these samples removed. The bacterial supply effect remained significant and the supply-by-bias interaction remained in the same direction, though it became marginal (*P* = 0.059), consistent with the reduced statistical power from excluding 35 samples (Table S1-2). Second, we verified the extent to which our experimental treatments modified the dispersal in patches supplied with bacteria relative to those without. For example, if treatments were successful, then bacteria supplied patches should be less likely to be selected for dispersal relative to patches with no supply under the dispersal bias treatment. To test this, we examined our records of which patches were selected for dispersal and measured differences in dispersal propensities (i.e., frequency that a patch type was selected for dispersal) between dispersal treatments and tested whether these differences differed by patch type. A GLM revealed marginally significant difference in the dispersal propensities between patch types (*P* = 0.053; Fig. S4B). Lastly, we confirmed that the position of microplate wells had no effect on yeast densities (*P* = 0.491; Fig. S5) by comparing the densities between wells within replicate after controlling for treatment effects via bootstrapping (see Supplementary Methods).

### Metapopulation model

We analysed a two-patch metapopulation model as a heuristic for understanding when dispersal bias amplifies an invader’s local benefit to a resident’s metapopulation size. A surprising result we found in our experiment was that local yeast (*M. reukaufii*) densities were positively correlated with local bacterial (*A. nectaris*) densities (Fig. 4), suggesting that bacteria can enhance local yeast productivity. As such, we represented the two effects of bacteria on a patch as separate levers, an increase in the local carrying capacity of bacteria-present patches (productivity effect) and a reduction in their per-capita dispersal rate relative to bacteria-absent patches (dispersal bias effect). We remain agnostic about the biological cause of the productivity effect, as our data cannot distinguish resource inputs, altered yeast metabolism, or other local mechanisms (see Discussion). With this model, we then asked how these two effects jointly shape yeast metapopulation size, and whether they reproduce the patterns observed in our experiment.

To simplify analysis, we did not explicitly model the densities of the bacterial species and simply assumed that its effects are reflected in the demographic and dispersal parameters of the resident yeast.

Our two-patch model takes the form:

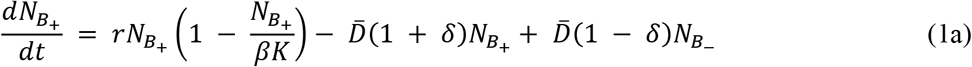

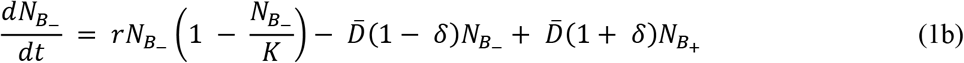

where 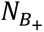 and 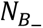 are the densities of the resident yeast in patch *B*_+_ (bacteria-present) and patch *B*_−_ (bacteria-absent), respectively. The parameters *r* and *K* represent the local intrinsic growth rate and baseline carrying capacity (*i.e.*, the carrying capacity in the absence of productivity enhancement by bacteria), while *D̅* denotes the average per-capita dispersal rate across the metapopulation. The parameters *β* and *δ* control asymmetries in local carrying capacity and per-capita dispersal rate between the two patch types. Specifically, *β* represents the local productivity enhancement factor induced by the presence of bacteria, whereas *δ* represents asymmetry in per-capita dispersal rates between patches induced by local bacterial pH modification and pollinator preference, with *δ* < 0 indicating a lower per-capita dispersal rate in patch *B*_+_ relative to patch *B*_−_ (*i.e.*, dispersal bias treatment). In the Appendix, we provide a detailed derivation of why pollinator patch preferences generate asymmetry in patch-specific per-capita dispersal rates (Equation A1 and A2). Importantly, *β* introduces asymmetry in carrying capacity between local patches while also increasing the total carrying capacity of the metapopulation, whereas *δ* modifies the asymmetry in per-capita dispersal rates between patches while holding average per-capita dispersal rate *D̅* fixed.

We analysed the model described by Equation 1 by numerically solving for its steady-state solution. We assume *r* = *K* = *D̅* = 1, which reduces the number of free parameters to two, corresponding to the two effects of bacteria in our experiment, the local productivity enhancement from the addition of bacterial cells (*β*) and dispersal bias (*δ*). With this reduced model, we computed the metapopulation size at varying levels of *β and δ* by solving for its steady-state solution numerically in R and taking the average of local equilibrium densities (denoted 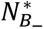 and 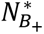) across both patches. To obtain a heuristic understanding of why metapopulation size varies at all, we derived an analytical condition under which metapopulation size is maximised in our model, as predicted by the theory of ideal free distribution (IFD; Fretwell & Lucas, 1969; Holt, 1985). In brief, IFD predicts that mean fitness (i.e., average per-capita growth rate excluding dispersal) is maximised when fitness does not vary across patches. At equilibrium, this corresponds to the condition where the density in each patch equals its local carrying capacity, since any deviation would imply unequal fitness: one patch would be underfilled (positive per-capita growth) while the other would be overfilled (negative per-capita growth). One trivial case in which this condition holds is when *D̅* = 0. However, since we assume dispersal (*D̅* = 1), the remaining condition that satisfies IFD occurs when the number of migrants in each patch cancels out, 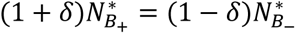 Substituting the local carrying capacities, *β* and 1, for the equilibrium densities 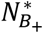 and 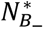 and re-arranging yields the condition *δ* = (1 − *β*)/(1 + *β*). We verify that this condition corresponds to the maximisation of metapopulation size by comparing it to simulations.

We note here that the two-patch model follows the same dispersal rule as the experiment but reduces it to a single patch of each type (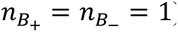). As a result, dispersal asymmetry between the two patch types depends solely on the preference differential (captured by *δ*), whereas in the experiment it also depends on the number of patches of each type, which the bacterial supply treatment sets (*i.e.*, 0, 3, or 6). In the Appendix, we provide a detailed account of how the treatments in our experiment map onto a general multi-patch metapopulation model and show that both the two-patch model and the eight-patch experiment are special cases of it. To ensure that the two-patch simplification results are robust to variation in the number of patches, we provide additional analyses of the general multi-patch model in the Appendix. For simplicity and clarity, we present only the results of the two-patch model and only refer to the Appendix for the general case.

## RESULTS

### Effects of bacterial supply, dispersal bias, and dispersal frequency on yeast metapopulation size

Bacterial supply significantly increased overall yeast metapopulation size (estimate = 0.028; 95% CI: 0.005 to 0.052; *χ*^2^ = 5.68, *P* = 0.017; Table 1; Figs. 2 & 3A), and the magnitude of this effect depended on dispersal bias (interaction estimate = 0.056; 95% CI: 0.010 to 0.102; *χ*^2^ = 5.74, *P* = 0.017; Table 1). Specifically, bacterial supply increased yeast metapopulation size under biased dispersal, but not under random dispersal (Fig. 3B). In contrast, neither dispersal frequency (estimate = 0.011; 95% CI-0.026 to 0.049; *χ*^2^ = 0.67, *P* = 0.414) nor dispersal bias (estimate = 0.068; 95% CI-0.045 to 0.180; *χ*^2^ = 2.02, *P* = 0.156) alone had a significant main effect on yeast metapopulation size (Table 1). The interaction between bacterial supply and dispersal frequency was marginally significant (estimate = 0.045; 95% CI: 0.000 to 0.091; *χ*^2^ = 3.76; *P* = 0.053; Table 1). The non-significant main effect of dispersal bias is expected, as the experimental design ensures that dispersal bias is only functionally consequential when bacterial sink populations are present (i.e., when bacterial supply > 0). Only under these conditions would pH values differ between patches and asymmetric dispersal be possible under dispersal bias. Thus, the non-significant effect of dispersal bias in isolation reflects a sensible experimental setup. Lastly, local yeast densities also positively correlated with local bacteria densities (*ρ* =0.140, *P*<0.001; Fig. 4), an unexpected result that suggests that bacteria have a net positive (rather than purely competitive) effect on yeast population size.

**Figure 2.**
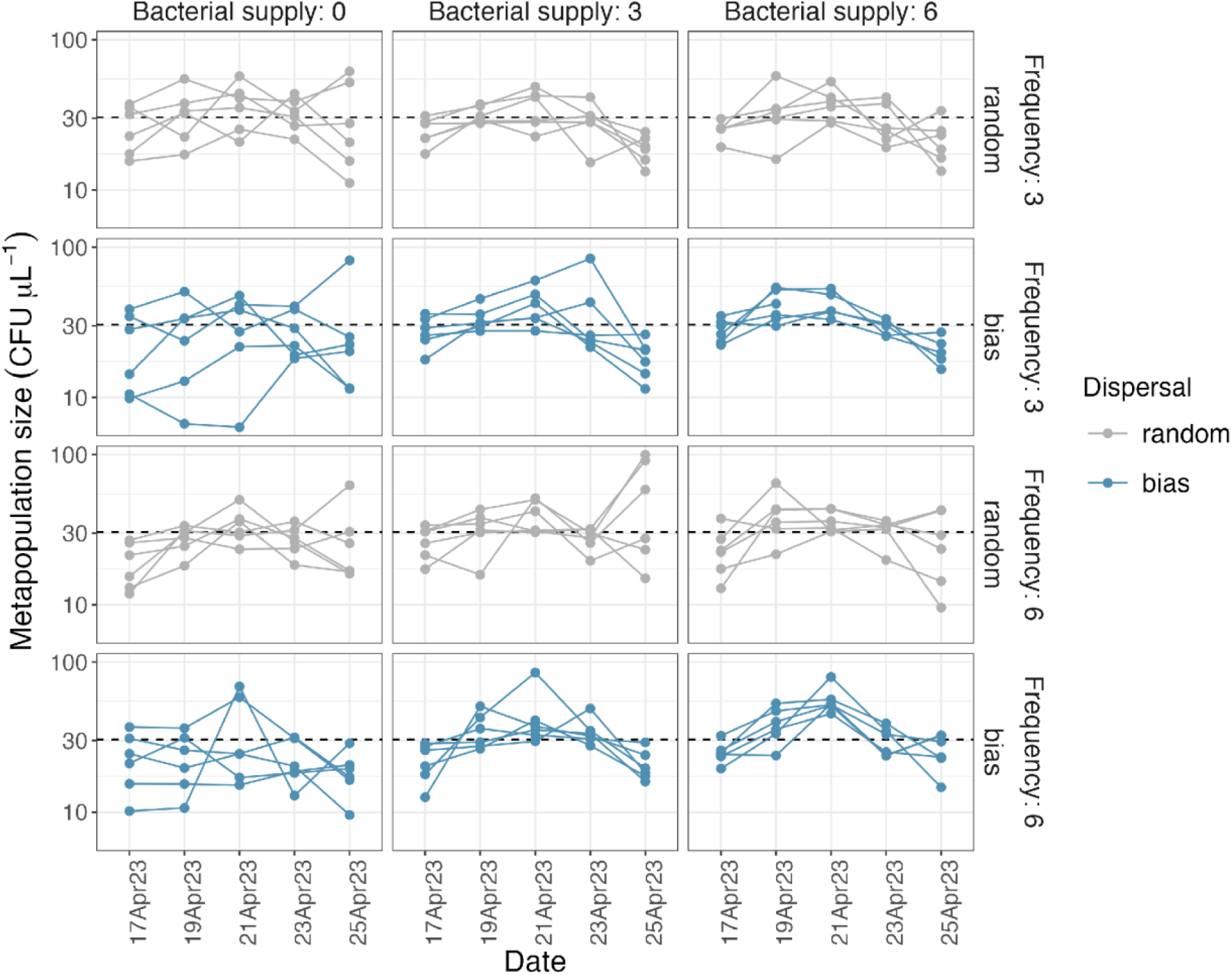
Time-series of yeast metapopulation size. Points connected by lines show the yeast metapopulation size (CFU µL^-1^) of a single replicate over time, with grey indicating random dispersal and blue indicating biased dispersal. Each facet corresponds to a treatment and its six replicate time series. The dashed horizontal line corresponds to the global mean (∼30.5 CFU µL^-1^).

**Figure 3.**
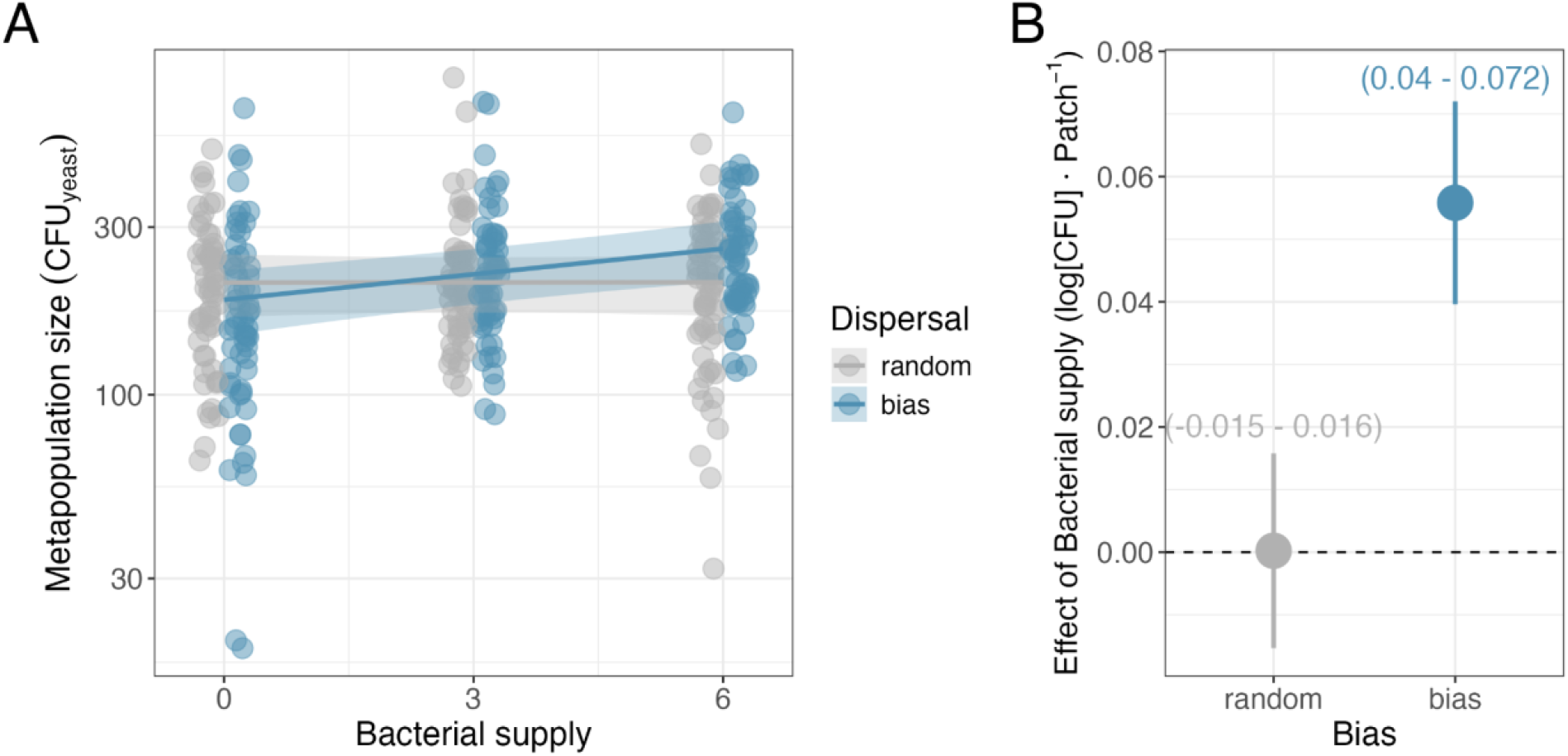
Effects of bacterial supply and dispersal bias on yeast metapopulation size. **(A)** Points show the yeast metapopulation size (CFU µL^-1^) of experimental replicates (n = 354) plotted against the bacterial supply treatment, with grey and blue indicating random and biased dispersal, respectively. Lines show predicted values of the fitted GLMM with shaded region indicating ±1 SE. (**B**) Points show the estimated marginal effect of bacterial supply on yeast metapopulation size across levels of dispersal bias, evaluated at the mean dispersal frequency (4.5) with error bars indicating ±1 SE.

### Theoretical predictions of metapopulation model

Theoretical predictions agree qualitatively with our experimental results that on average, local enhancement of productivity increases regional metapopulation size and that this effect is greater when the more productive patch has relatively lower per-capita dispersal rates (Fig. 5). Note that because we assume *r* = *K* = *D̅* = 1 and 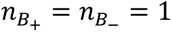 for analytical tractability, our comparisons between theoretical predictions and experimental results are based on the relative sign of effects rather than their exact numerical values. Specifically, our theoretical model predicts that increasing the productivity of the bacteria-present patch (*β* > 1) increases metapopulation size (Fig. 5), consistent with how bacterial supply increased yeast metapopulation size by ∼3% per patch supplied in the experiment (Table 1; Fig. 3B). Furthermore, the effect of lowering per-capita dispersal rate in the bacteria-containing patch relative to the bacteria-absent patch (*i.e.*, decreasing *δ*, analogous to dispersal bias treatment) reduced metapopulation size when carrying capacities were similar, but increased it as the carrying capacity of the bacteria-containing patch increased (*β* ≫ 1; Fig. 5A). In other words, dispersal bias increases metapopulation size only when bacteria have a strong enough local positive effect on yeast, consistent with the significant interaction term between bacterial supply and dispersal bias observed experimentally (predicted lines cross in Fig. 3A). Furthermore, we confirmed that these qualitative results hold in a metapopulation with eight patches, although effect sizes diminish as the number of patches increases (Fig. A2). We discuss this limitation further in the Discussion. Lastly, we found that metapopulation size is maximised when the total number of migrants in each patch is equalised, in accordance with the ideal free distribution (unpacked in Discussion; Fig. 5B).

**Figure 4.**
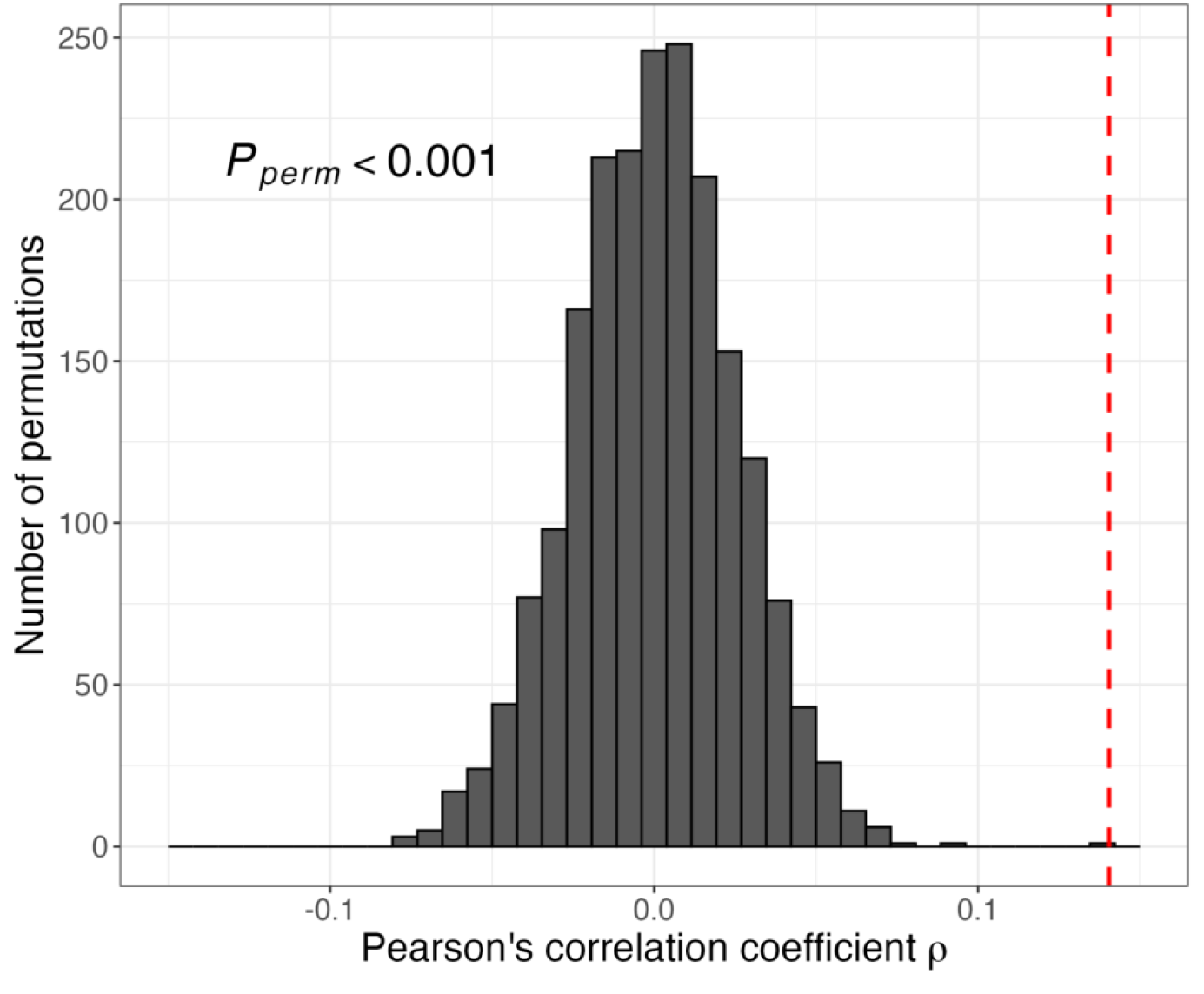
Permutation test for the association between local densities of *M. reukaufii* and *A. nectaris* (n = 2,832). The histogram shows the null distribution of Pearson correlation coefficients generated from 2,000 block-structured permutations to account for statistical non-independence (see Methods). The observed coefficient (*ρ* = 0.140, dashed vertical line) fell above the entire null distribution (*P* < 0.001).

**Figure 5.**
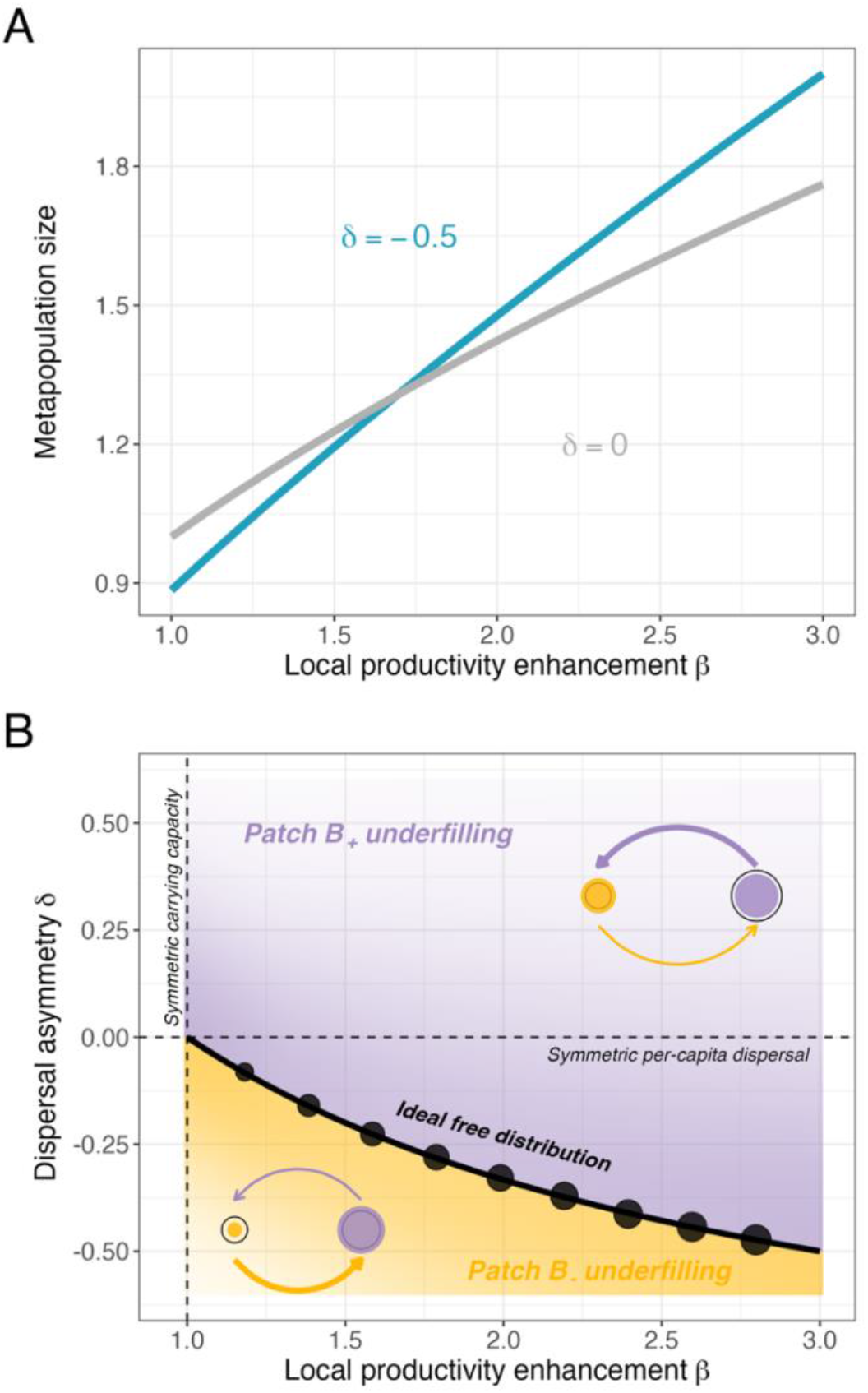
Theoretical predictions of metapopulation size with asymmetric per-capita dispersal rates and carrying capacities. (**A**) Metapopulation size increases with productivity enhancement (*β*) and increases more steeply when patch *B*_+_ has a lower per-capita dispersal relative to *B*_−_ (blue curve relative to grey curve). (**B**) As *β* increases, a stronger dispersal bias (*δ* < 0) is needed to maximise metapopulation size (decreasing curve). The curve satisfies 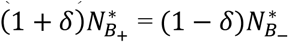 and corresponds to the ideal free distribution where both patches send equal numbers of migrants at equilibrium; above and below it, patch *B*_+_ and *B*_−_, respectively, contribute more migrants. Numerical simulations confirm that this curve traces the *δ* that maximises metapopulation size as function of *β* (black points, sized in proportion to metapopulation size). The stock-and-flow diagram illustrates the asymmetry in carrying capacity (open circle size) and per-capita dispersal rate (arrow thickness), and the resulting under- and over-filling (coloured circle size relative to open circle). Dashed vertical and horizontal lines indicate symmetric carrying capacity and per-capita dispersal. In both panels, *r* = *K* = *D̅* = 1.

## DISCUSSION

Our study demonstrated that transient invaders may affect a resident species by modifying both local habitat conditions and regional connectivity, despite being unable to persist in the absence of constant external immigration. In particular, we found that addition of an externally-sustained bacterial species increased the resident yeast metapopulation size, and that this effect was larger when they also reduced dispersal rates in the patches in which they occurred. Furthermore, our theoretical results align with experiments, and suggest that the mechanism by which dispersal bias increases yeast metapopulation size is through equalising per-capita growth rates across patches, thereby bringing the metapopulation size closer to the theoretical maximum defined by the ideal free distribution. Taken together, these results demonstrate that transient invaders can impose demographic consequences on resident species through the interaction between local and regional processes. Below, we connect our results to metapopulation theory, discuss the biological mechanisms that may have generated these patterns, place our findings within broader ecological frameworks, and discuss key study limitations as well as opportunities for future research.

### Dispersal and metapopulation size

Our study provides a rare empirical demonstration of how biotically-induced dispersal asymmetries can alter metapopulation size. Furthermore, the analysis of our theoretical model connects our experimental results to well-known effects of dispersal in spatially heterogeneous landscapes. In particular, theory shows that when carrying capacities vary spatially, dispersal typically reduces metapopulation size because productive patches export more migrants than they receive, causing greater overfilling of low-capacity patches than underfilling of high-capacity patches and thus stronger overall density dependence (Holt, 1985; McPeek & Holt, 1992; Pacala & Roughgarden, 1982). This demographic cost of dispersal underlies the “slower disperser wins” phenomenon (Dockery et al., 1998; Hastings, 1983) and explains how a locally inferior competitor can coexist with a competitively superior species with faster dispersal rates (Abrams & Wilson, 2004; Amarasekare & Nisbet, 2001; Pacala & Roughgarden, 1982). With respect to our experiment, the demographic cost of dispersal in a landscape with heterogeneous carrying capacities provides a plausible explanation for why the positive effect of bacterial supply on metapopulation size was absent when dispersal was random (Fig. 3).

In agreement with McPeek & Holt (1992), our study shows that the detriments of dispersal on metapopulation size can be nullified if differences in local per-capita dispersal rates offset differences in local carrying capacities. The degree of this offset can be interpreted as a deviation from the ideal free distribution, the condition under which local growth rates are equalised and metapopulation size is maximised (Fig. 5B). Our formulation of dispersal slightly differs from that of McPeek & Holt (1992); rather than assigning each patch an independent dispersal parameter, our model contains a single parameter that controls the asymmetry in per-capita dispersal between patches. This formulation reflects dispersal asymmetry arising from the foraging preferences of the dispersal vector (i.e., pollinators) as opposed to individual microbes moving between flowers, such that a single parameter captures a metapopulation-level process that generates patch-specific differences in per-capita dispersal rates. More broadly, our results also align with empirical studies showing that movement between habitats can modify demographic parameters (Donahue et al., 2003; Fox, 2007), while extending this idea to the modification of dispersal parameters themselves.

### Biological mechanisms: Why does bacterial supply increase yeast densities?

Our experimental results appear to contradict previous work showing that the reduction in pH by bacteria inhibits yeast growth (Chappell et al., 2022; Figs. 3-4 & S1). We posit that experimental differences may have contributed to the apparent disagreements. In the present experiment, densities of yeast and bacteria were approximately the same at the beginning of each dilution cycle (∼100-200 cells/µL; Fig. 1, see Methods). Thus, nectar acidification by the bacteria occurred concurrently with yeast population growth, possibly preventing pH from dropping fast enough to suppress yeast growth. However, this explanation is only partial because it does not explain the observed positive effect of bacterial supply on yeast densities. We posit that the addition of bacterial cells may have also introduced resource subsidies that enabled yeast to reach higher densities. Although each inoculation event contributed only ∼1% of the nectar volume, additional inputs may have arisen from metabolites released by bacterial cells or from cell lysis, beyond those present in the nectar medium used to suspend cells prior to inoculation. While this mechanism has not been confirmed experimentally, it is consistent with reports of *Acinetobacter* spp. releasing metabolic by-products in low-nitrogen nectar environments (Álvarez-Pérez et al., 2019; Kato et al., 2025; Morales-Poole et al., 2023)

Alternatively, bacteria may have increased yeast productivity by reprogramming yeast metabolic pathways, either through induced plasticity or evolutionary change. Plastic metabolic responses have been documented in other fungi whose gene expression and metabolism respond to ambient pH (Peñalva & Arst, 2004; Jouhten et al., 2016), but such responses are likely species-specific, and whether this holds for *M. reukaufii* is unknown. Rapid evolution has likewise been documented in *M. reukaufii* (∼50 transfers; Chappell et al., 2022), but with only four growth-dilution cycles it is unlikely to explain the pattern in our experiment. Taking these possibilities together, we suspect that resource subsidies from bacterial metabolites or cell lysis remain the most parsimonious explanation for the increased yeast productivity, which is plausible given that many microbes secrete metabolites (Douglas, 2020) and that dead cells can affect local communities (Hao et al., 2026). Distinguishing these possibilities will require directly manipulating nectar chemistry rather than indirect inference via correlations, and identifying the biochemical pathways involved remains an important direction for future work.

### Meta-ecosystems and nectar microbiome

The increased yeast productivity we found may be specific to our experimental conditions and model organisms, but the process may be more general and relevant across ecosystems. The particular finding aligns with core ideas in meta-ecosystem ecology, which emphasizes how flows of matter and energy shape ecosystem dynamics (Gounand et al., 2018; Loreau et al., 2003; Massol et al., 2017; Polis et al., 1997). By focusing on connections across ecosystem boundaries, this framework considers a broader set of spatial flows than traditional metapopulation approaches would, including the movement of inorganic nutrients, organic matter, and other non-living biotic materials in addition to living organisms. In nectar microbes, for instance, floral nectar represents a primary habitat but not the only one they occupy. As nectar yeasts may overwinter in bee nests (Pozo et al., 2018) or use soil and plant surfaces as refugia, their re-colonisation of nectar during the flowering season (Vannette, 2020) may serve as an ecological corridor, allowing materials to flow between these otherwise distinct ecosystems and habitat. Together, these examples highlight that dispersal moves not only organisms but also the energy and nutrients associated with them from previous places, prompting questions about the extent to which these cross-ecosystem fluxes affect local microbial communities.

### Experimental caveats and theoretical assumptions

Our experimental and theoretical approaches necessarily involve several caveats and simplifying assumptions. For tractability, our two-patch model formalism ignores dispersal among identical habitats, implicitly assuming that such movements have negligible effects on metapopulation dynamics. While this may be a reasonable assumption within an idealised experimental setting like ours, movement between identical habitats is likely to be important in real-world systems with extinction-recolonisation dynamics (*e.g.*, Brown & Kodric-Brown, 1977). Although the general multi-patch model includes within-type dispersal, incorporating disturbances and stochastic fluctuations would be a necessary next step to examine how these asymmetries in dispersal and productivity may interact with extinction-recolonisation dynamics. Moreover, our model also excludes explicit dynamics of the bacteria species which we consider justifiable given that they only persist through external supply much like an abiotic resource in a chemostat. Adding explicit species interactions would be straightforward to implement, by indexing parameters by patch and species, as would stochasticity, for which our simulation code already provides as arguments. However, these extensions would expand the parameter space combinatorially and are beyond the scope of the present study, so we leave them for future work.

One notable simplification, in both the theory and experiment, is that the benefit of dispersal bias depends on the number of patches in the metapopulation. As we show in the Appendix, the dispersal asymmetry generated by pollinator preferences arises because a donor patch cannot also be its own recipient, so its weight is removed from the pool of possible destinations (a recipient normalisation; Equation A2). This makes emigration and immigration asymmetric, and the asymmetry weakens as patches are added, since removing any single patch matters less when more remain, vanishing altogether in the limit of infinitely many patches. We term this a “finite-size effect”. Reassuringly, our analysis of the general multi-patch model shows that it does not undermine our conclusions, as the asymmetry, and hence the benefit of dispersal bias, remains relevant across the range of patch numbers including those used in our experiment (Appendix; Fig. A2).

Beyond our idealised system, this finite-size effect may also be biologically realistic. Because nectar microbes disperse only by hitchhiking on pollinators, the ecologically relevant number of patches is not the total number of flowers in a landscape but the set of flowers a single pollinator visits and holds in memory. Because hummingbirds are the primary pollinators of *Diplacus* (Fetscher & Kohn, 1999; Fetscher et al., 2002; Streisfeld & Kohn, 2007), known for their territoriality and trap-lining foraging behaviour (Stiles, 1975; Gill, 1988), individuals will likely sample a limited set of flowers with little overlap between birds. Thus, the number of patches governing dispersal asymmetry should be much closer to this small per-pollinator neighbourhood than to the landscape total, placing this natural system in the regime where the finite-size effect operates.

Our results should be interpreted considering how dispersal was experimentally implemented, specifically where bacteria were placed and the realised dispersal pattern generated by the sampling algorithm. Because bacteria were always inoculated into the same patches every cycle, our results could in principle reflect spatial placement (e.g., edge effects on the PCR plate) rather than treatment *per se*. However, we found no evidence for such location-specific effects on yeast densities (Supplementary methods; Fig S5). As for the realised dispersal pattern, the dispersal bias treatment produced only a marginally significant difference in dispersal propensities between patch types (bacteria-absent vs. bacteria-present; *P* = 0.053; Fig. S4), yet even this small asymmetry amplified the positive effect of bacterial supply. While this is a genuine limitation, it would be more concerning if our predictions rested on a main effect of dispersal bias. Instead, both our experiment and theoretical model treat dispersal bias as a modulator of the effect of bacterial supply, so its biological effect size depends jointly on the magnitude of the dispersal bias and the strength of the local bacterial effect.

## Conclusion

This study experimentally demonstrated that a species invading a sink habitat can influence a resident metapopulation by modifying local conditions and altering metapopulation connectivity. More broadly, our results highlight that the movement of organisms can itself generate the spatial variation in carrying capacity that determines the metapopulation consequences of dispersal. Biologically, this is most likely when dispersing organisms carry resources acquired in previous habitats. Our experiments thus demonstrate the possibility that transient invaders may still leave observable ecological effects even when the habitat modifying individuals cannot survive in the habitat. Given the prevalence of sink populations (Dias, 1996; Pulliam, 1988) in general and of unsuccessful microbial invasions in particular (Amor et al., 2020; Mallon et al., 2018), we suggest that it is worth investigating the extent to which these transient invaders structure local communities and metapopulations in natural ecosystems.

## Appendix

Here we formalise the dispersal rule used in our experiment as a general multi-patch metapopulation model, in which the number of patches, the number of patch types, and their composition may each take any positive integer value. This serves two purposes. First, it shows that both the two-patch model of the main text and the eight-patch experiment are special cases of a single model, differing only in the number and composition of patches rather than in the underlying dispersal process. Second, by analysing the general model we test whether the qualitative insights from the two-patch model still hold in the multi-patch regime of our experiment. Together, these establish that the difference in patch number between model and experiment does not reflect a difference in mechanism.

## General multi-patch dispersal model

The general model describes a single resident species (*e.g.*, yeast) occupying *M* patches belonging to *L* types, where the number of patches *M*, the number of types *L*, and the number of patches of each type *n_ℓ_* (with 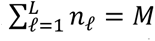) may all take any positive integer value. The two-patch model of the main text (*M* = 2, *L* = 2, one patch per type) and our eight-patch experiment (*M* = 8, *L* = 2) are both recovered by fixing these quantities. In the following, we assume the case of *L* = 2, as in our system. Each patch *i* ∈ {1, …, *M*} has density *N_i_*, intrinsic growth rate *r_i_*, local carrying capacity *K_i_*, and pollinator-preference weight *w_i_* (set by nectar pH in the experiment). These patch-specific parameters *r_i_*, *K_i_*, *w_i_* are determined by the patch’s type. With *L* = 2, patches can be one of two types: *B*_−_ or *B*_+_ for bacteria-absent and bacteria-present, respectively (*i.e.,* ℓ ∈ {*B*_−_, *B*_+_}) and dispersal is governed by a single metapopulation-wide rate constant *D̅*, so differences in patch-specific dispersal rates arise solely through preference weights *w_i_*. Biologically, this means that the number of pollinator visitations is fixed across the metapopulation (“dispersal frequency” in our experiment) but some flowers (*i.e.*, patches) are visited more often than others whenever pollinators have preferences (*i.e.*, 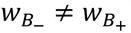).

Dispersal follows the same rule as the experiment in which each dispersal event draws a donor patch with probability proportional to its weight, then a recipient among the remaining patches, again proportional to weight. Because there are a finite number of patches, the probability of drawing a *donor* of type ℓ is proportional to the number of type-l patches multiplied by their weight, *n*_ℓ_ *w*_ℓ_. The *recipient* is drawn the same way, but only from the remaining patches not selected as a donor, so the probability that the recipient is of type ℓ is proportional to the weight of the type-l patches still available, divided by the total weight of those remaining patches. Writing 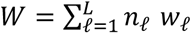 for the total weight, the dynamics of patch *i* are

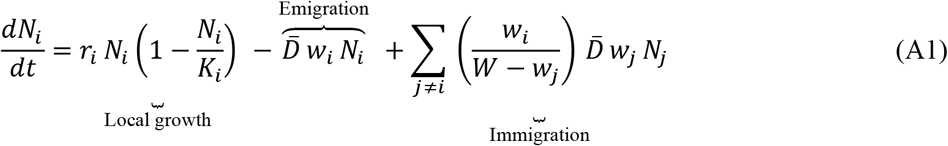

Emigration out of patch *i* scales with its preference weight *w_i_* while immigration is the sum, over every other patch *j*, of the emigrant pool leaving *j* (*i.e.*, *D̅ w_j_ N_j_*) times the proportion of the remaining weight that patch *i* contributes to the total weight after removing the donor 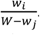. The recipient normalisation

*W* − *w_j_* excludes the donor patch *j* from the pool of possible destinations, and this donor-exclusion term is the origin of the between-type dispersal asymmetry (see *Consequences and scope*).

### Reduction to the two-patch model presented in the main text

With one patch of each type (*i.e.*, 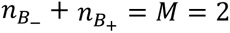), a donor has only one possible recipient, so every migrant leaving one patch enters the other. The emigration and immigration terms of patch *i* then simplify (using *W* − *w_j_* = *w_i_*) to

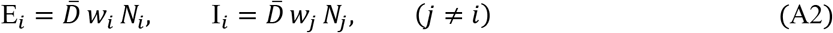

Assuming spatially-homogeneous intrinsic growth *r*, the general model at *M* = 2 reduces exactly to the two-patch model of the main-text, with the dispersal and local carrying capacity asymmetry parameters, *δ* and *β*, emerging from local patch weights and carrying capacities respectively:

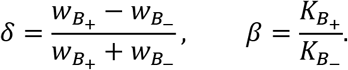

Here, *δ* < 0 occurs when pollinators prefer *B*_−_ patches (*i.e.*, 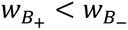) and corresponds to the dispersal bias treatment of the experiment, while the local productivity-enhancing effect of bacteria gives *β* > 1. Thus, the two-patch model follows the exact same *dispersal rule* as the experiment and differs only in parameter values. However, when the types contain unequal numbers of patches (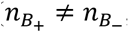, or more generally when the *n*_ℓ_ are not all equal), the counts no longer cancel and dispersal rate of each type becomes proportional to *n*_ℓ_*w*_ℓ_ and not on the weights alone. We develop the consequences of this in the following section.

#### Consequences and scope

The two-patch model is not a separate model but a limiting case of Equation A1, with *M* = 2 and one patch of each type (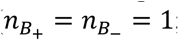; Fig. A1). Two features separate this regime from the eight-patch experiment, and both concern how the between-type dispersal asymmetry is set.

The first is that the type counts need not be equal. Each type sends out migrants at a rate proportional to *n*_ℓ_ *w*_ℓ_, so when 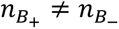 the between-type asymmetry is shaped by both the counts *n*_ℓ_ and the weights *w*_l_. In the balanced two-patch case the counts cancel and only the weights matter, as captured by *δ*. In the experiment, patches with and without bacteria are never present in equal numbers, because the bacterial-supply treatment fixes the number of bacteria-present patches at 0, 3, or 6 of the 8 (so 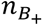 is never 4). This imbalance means that supply and bias both act on the between-type flow, and they act in opposite directions. Lower weight on *B*_+_ (the bias treatment, *δ* < 0) directs migrants from *B*_−_ to *B*_+_, whereas adding *B*_+_ patches (the supply treatment) directs migrants from *B*_+_ to *B*_−_. Supply therefore works against bias, so the benefit of dispersal bias is unimodal in the number of bacteria patches. It vanishes at both extremes, where a single patch type makes bias degenerate, and peaks at an intermediate composition (Fig. A2B). The peak sits close to an even split of the two types but slightly below it, and shifts further below the centre as the productivity gap *β* widens. The reason is that the benefit of bias is the rescue of productive *B*_+_ patches from being drained into the low-productivity *B*_−_ patches. Each *B*_+_ patch is rescued more when it is rare and surrounded by many draining (relative) sinks, so the metapopulation gains most when the productive patches are a minority. The larger their productivity advantage *β*, the more value concentrates in those few patches, and the more the optimal benefit skews toward fewer patches.

The second is that the asymmetry weakens as the number of metapopulation patches increases. This weakening effect originates from the donor-exclusion term *W* − *w_j_*, which is present because a patch cannot disperse to itself. As patches are added at fixed composition (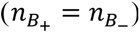), each contributes a vanishing share of the total weight (*w_j_*/*W* ≈ 0), such that *W* − *w_j_* ≈ *W* and the donor-exclusion asymmetry washes out, which we describe as a “finite-size effect” (Fig. A2A).

Taken together, both features point to the two-patch model as the regime where the beneficial effect of dispersal bias is exaggerated. The benefit grows as the metapopulation shrinks (the finite-size effect), and at the smallest size that contains both types, *M* = 2, the two types are necessarily balanced (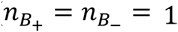), which is close to the composition that maximises the benefit at any *M*. To confirm that the analytical insights derived from the two-patch model hold under the parameter regime that aligns with our experiment, we numerically solved for the metapopulation equilibrium of Equation A1 across the experimental parameter range and present these results below (Fig. A2).

## Results

The results show that biased dispersal remains more beneficial than random dispersal whenever bacteria enhance local productivity, even when the composition of patch type varies (due to the bacterial supply treatment) and when the number of metapopulation patches increases (Fig. A2). However, adding more patches and unbalancing the counts does indeed weaken the signal but the mechanism, deviation from ideal free distribution, and the sign of effects still hold. In particular, the benefits of dispersal bias remain observable within the parameter regimes of our experiment (dashed vertical lines; Fig. A2B), given that bacterial addition increases local productivity (*β* > 1).

**Figure A1.**
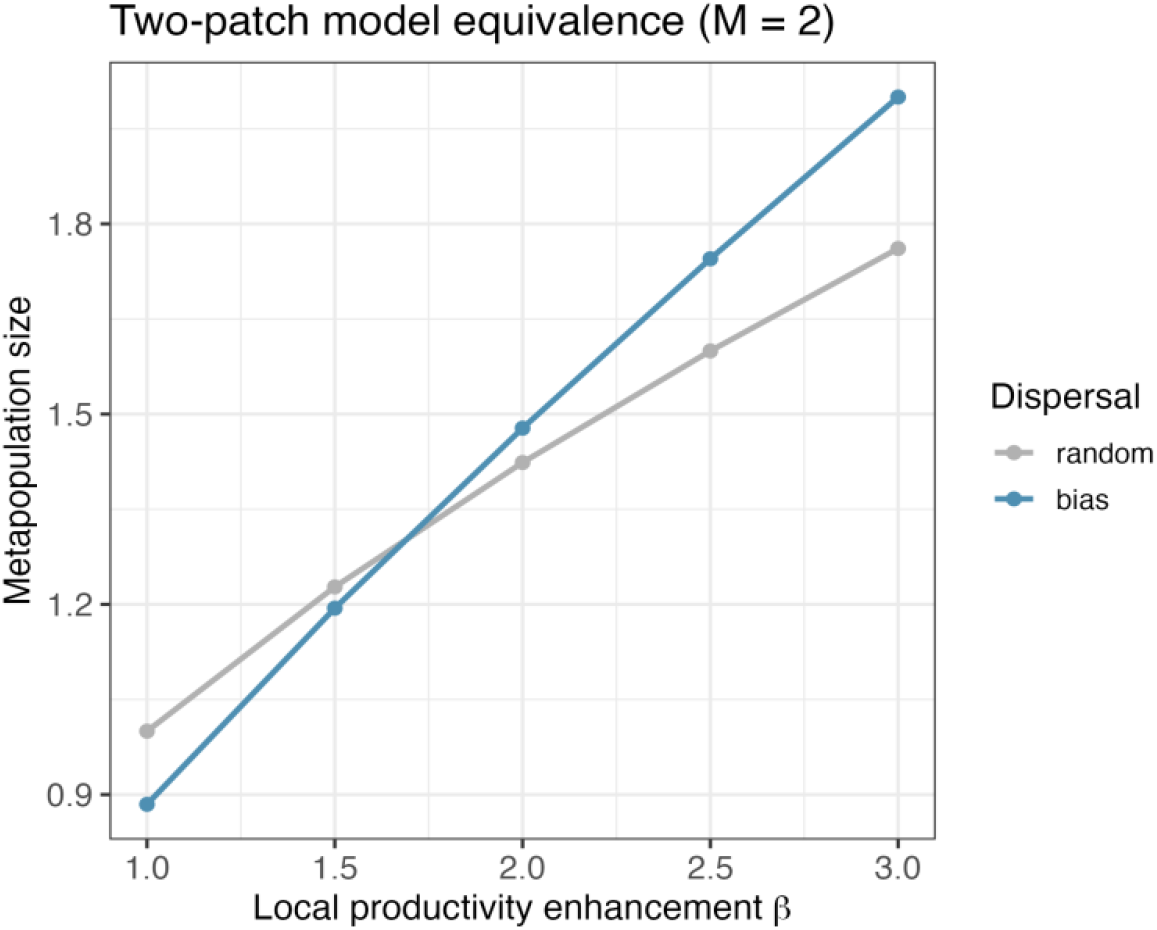
General multi-patch model is equivalent to two-patch model at M = 2. Plot shows the equilibrium metapopulation size as a function of local productivity enhancement *β* by bacteria. This figure reproduces Fig. 5A exactly using the general multi-patch model. The dispersal asymmetry parameter *δ* was set to 0 and -0.5 for random and biased dispersal respectively. The remaining parameters were *r* = 1, *K* = 1, and *D̅* = 1.

**Figure A2.**
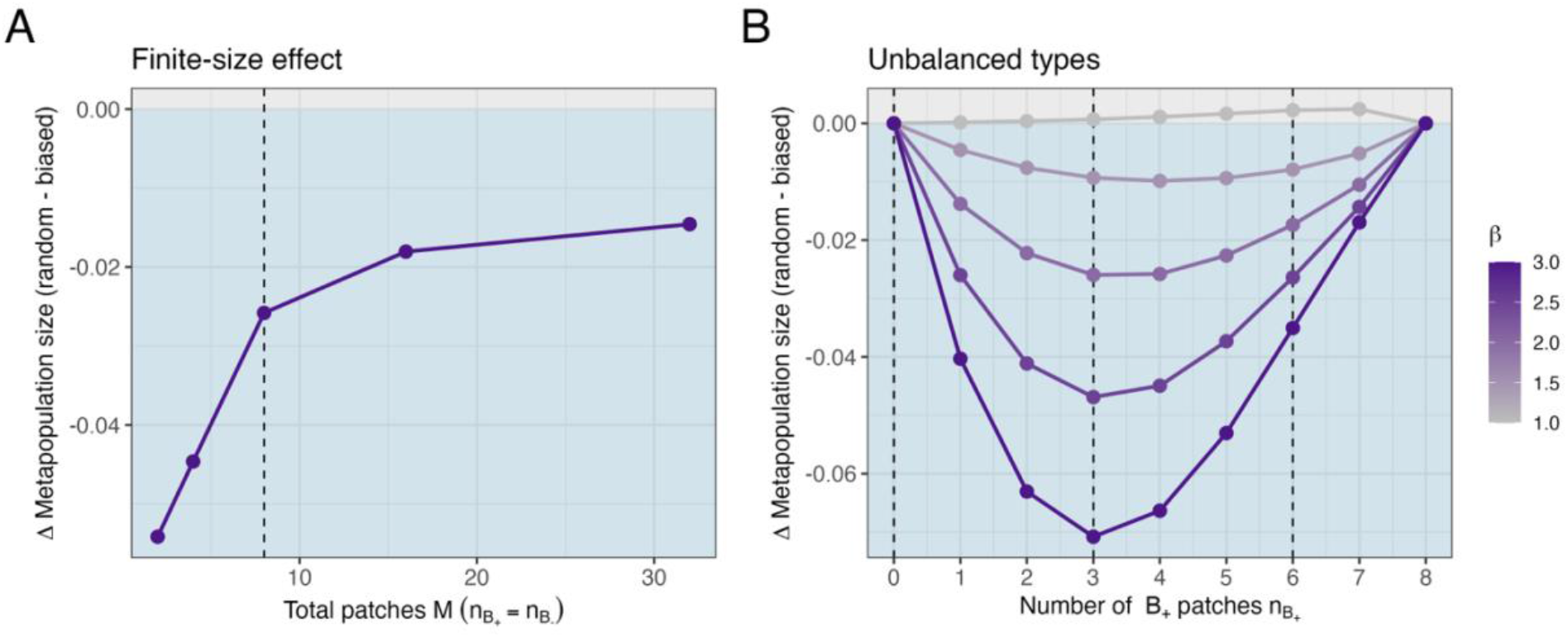
Effects of the number of metapopulation patches and unequal number of types. The y-axis shows the difference in equilibrium metapopulation size between random and biased dispersal. Negative values (blue region) indicate that biased dispersal yields a larger metapopulation relative to random dispersal. (**A**) At fixed composition and *β* = 2, the benefit of biased dispersal diminishes as the number of patches increases but remains present at the experimental size (*M* = 8; dashed line). (**B**) With the number of patches fixed at *M* = 8, the benefit of biased dispersal follows a unimodal curve in the number of bacteria-present patches 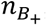, peaking at an intermediate, near-balanced composition (near 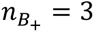) that shifts below an even split as the productivity enhancement *β* increases. Coloured lines and points show variation in local productivity enhancement by bacteria *β*, while dashed lines indicate parameters that correspond to the experimental design. Other parameter values are fixed at *r* = 1, *K* = 1, *D̅* = 1, and *δ* = −0.5 for biased dispersal.

