## Supplementary material for "Transient species can increase resident metapopulation size by modifying local conditions and regional connectivity"

#### Supplemental figures and tables

**Table S1. Forward model selection of dispersion terms**. ΔAIC and P (likelihood-ratio $\chi^2$ test) for terms added sequentially to the dispersion component of the negative binomial GLMM, each relative to the currently selected model. Columns show the result for the full (n = 354) and the reduced data set (n = 319) with uncountable local populations removed. Bacterial supply was retained for the full data, so steps 2 and 3 are tested on top of it; no term was retained for the reduced data. Replicate and sampling day were retained as random intercepts in the dispersion component of every model, including the baseline, and terms were kept when $P < 0.05$.

| Step | Term | $\Delta AIC_{full}$ | $P_{full}$ | $\Delta AIC_{reduced}$ | $P_{reduced}$ |
|---|---|---|---|---|---|
| 0 | Intercept-only | 0 | | 0 | |
| 1 | Bacterial supply | **-1.92** | **0.048** | -0.384 | 0.123 |
| 2 | Bias | 1.464 | 0.464 | -0.103 | 0.147 |
| 3 | Frequency | 1.048 | 0.329 | 1.894 | 0.745 |

**Table S2. Treatment effects on metapopulation size with uncountable local populations removed.** The model of Table 1 refitted to the reduced data set (n = 319; see Table S1). Bacterial supply was not retained in the dispersion component for these data, so only the conditional component is shown.

| Component | Term | $\chi^2$ | P |
|---|---|---|---|
| **conditional** | **Supply** | **6.067** | **0.014** |
| conditional | Bias | 0.707 | 0.401 |
| conditional | Frequency | 0.007 | 0.933 |
| conditional | Supply:Bias | 3.579 | 0.059 |
| conditional | Supply:Frequency | 2.051 | 0.152 |
| conditional | Bias:Frequency | 0.576 | 0.448 |

**Table S3. Continuous versus categorical parameterisation of bacterial supply.** Type II Wald $\chi^2$ ($P$) for the metapopulation-size GLMM with bacterial supply coded as a continuous predictor versus a three-level factor. Model comparison supports the continuous parameterisation (ΔAIC = -6.159). Factor-model terms involving bacterial supply have 2 degrees of freedom; all others have 1.

| Component | Term | $\chi^2$ -continuous ($P$) | $\chi^2$ -factor ($P$) |
|---|---|---|---|
| **conditional** | **Supply** | **5.681 (0.017)** | **6.418 (0.040)** |
| conditional | Bias | 2.016 (0.156) | 1.935 (0.164) |
| conditional | Frequency | 0.669 (0.414) | 0.707 (0.401) |
| **conditional** | **Supply:Bias** | **5.744 (0.017)** | 5.921 (0.052) |
| conditional | Supply:Frequency | 3.760 (0.053) | 4.797 (0.091) |
| conditional | Bias:Frequency | 1.113 (0.291) | 1.178 (0.278) |
| dispersion | **Supply** | **3.882 (0.049)** | 3.827 (0.148) |

**Table S4. Temporal robustness of treatment effects.** Type II Wald $\chi^2$ $P$-values for the metapopulation-size GLMM fit to the full time series and to each sampling day separately. Bold indicates $P < 0.05$.

| Term | Full series (n=354) | Day 2 (n=71) | Day 4 (n=70) | Day 6 (n=71) | Day 8 (n=71) | Day 10 (n=71) |
|---|---|---|---|---|---|---|
| Supply | **0.017** | 0.464 | **0.003** | **0.017** | 0.603 | 0.315 |
| Dispersal bias | 0.156 | **0.049** | 0.440 | **0.003** | 0.497 | 0.236 |
| Dispersal frequency | 0.414 | 0.362 | 0.220 | **0.012** | 0.862 | **0.042** |
| Supply:Bias | **0.017** | 0.273 | **0.013** | **0.006** | 0.257 | 0.202 |
| Supply:Frequency | 0.053 | 0.618 | 0.463 | 0.378 | 0.069 | 0.293 |
| Bias:Frequency | 0.291 | 0.079 | 0.399 | 0.913 | 0.635 | 0.216 |

**Table S5. Samples excluded owing to pipetting errors.** Each row is a metapopulation (bacterial supply, dispersal bias, dispersal frequency, replicate) on the sampling day for which one or more patches were unmeasured (recorded as NA in the raw data). Six such incomplete samples were excluded, leaving $n$ = 354 for analysis.

| Bacterial supply | Dispersal bias | Dispersal frequency | Replicate | Sampling day |
|---|---|---|---|---|
| 0 | biased | 3 | i | 2 |
| 6 | random | 3 | i | 4 |
| 3 | biased | 3 | vi | 4 |
| 6 | biased | 3 | ii | 6 |
| 6 | biased | 3 | ii | 8 |
| 6 | biased | 3 | ii | 10 |

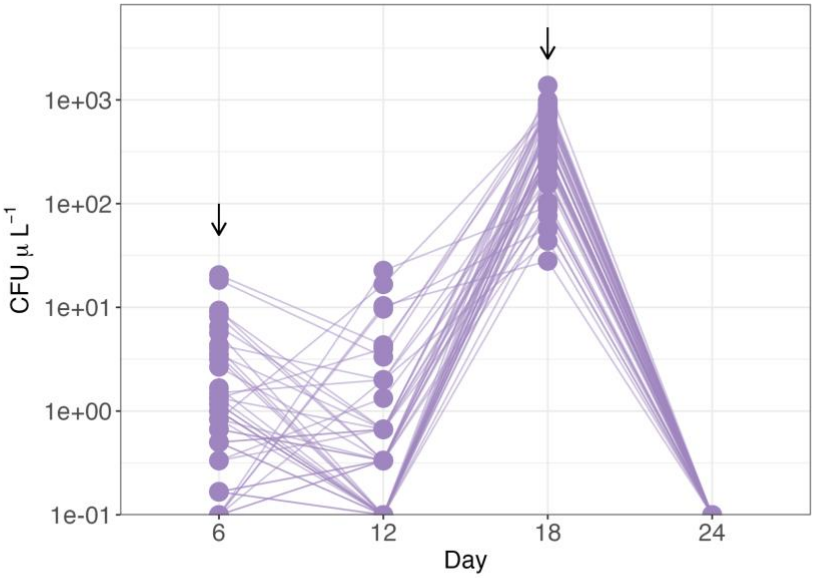


**Figure S1. Evidence of bacterial sink habitat from pilot study.** Points indicate the metapopulation-level bacteria density (CFU µL$^{-1}$) of a pilot experiment. Arrows show the time additional bacterial cells were supplied into the system from an external pool.

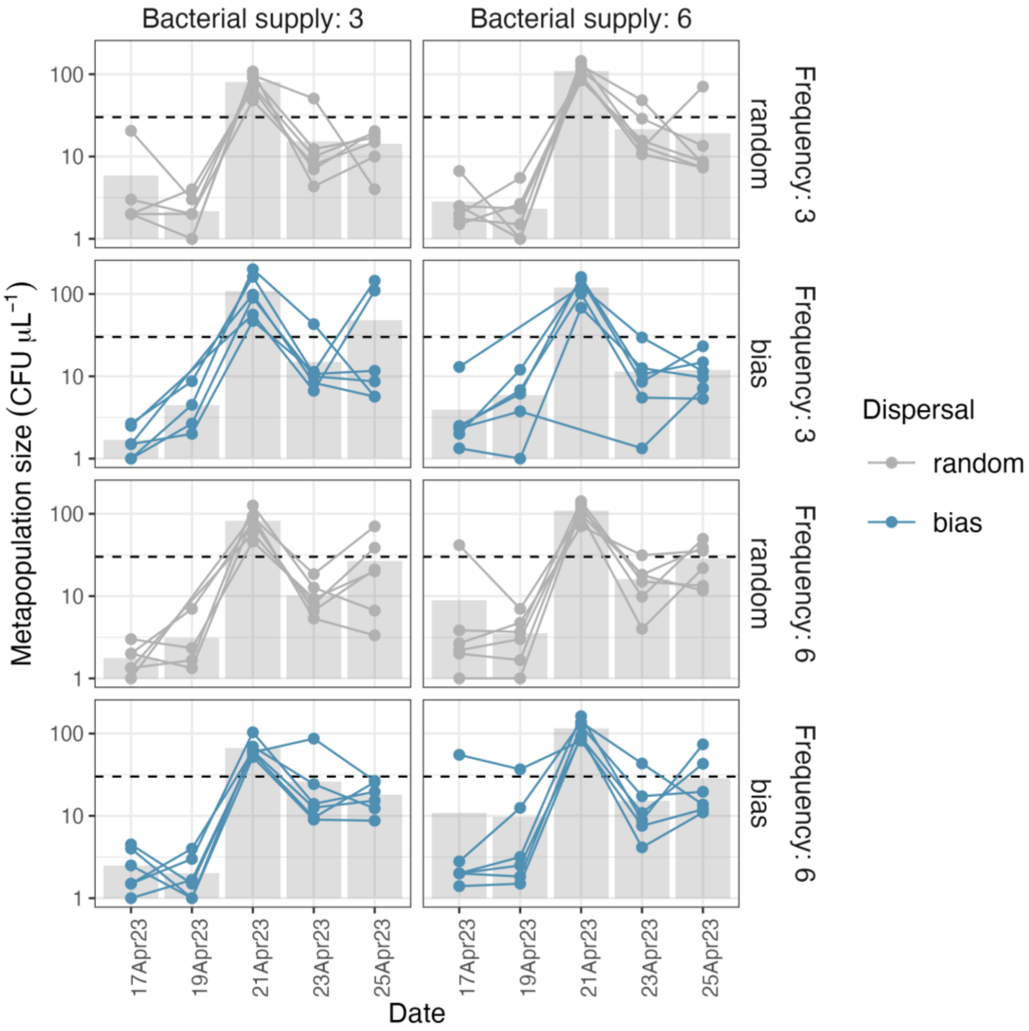


**Figure S2. Time-series of bacteria metapopulation size.** Points connected by lines show the bacteria metapopulation size (CFU μL$^{-1}$) of a single replicate over time, with grey indicating random dispersal and blue indicating biased dispersal. Each facet corresponds to a treatment and its six replicate time series. Gray bars indicate the average across replicates and the dashed horizontal line corresponds to the global mean (~30.0 CFU μL$^{-1}$).

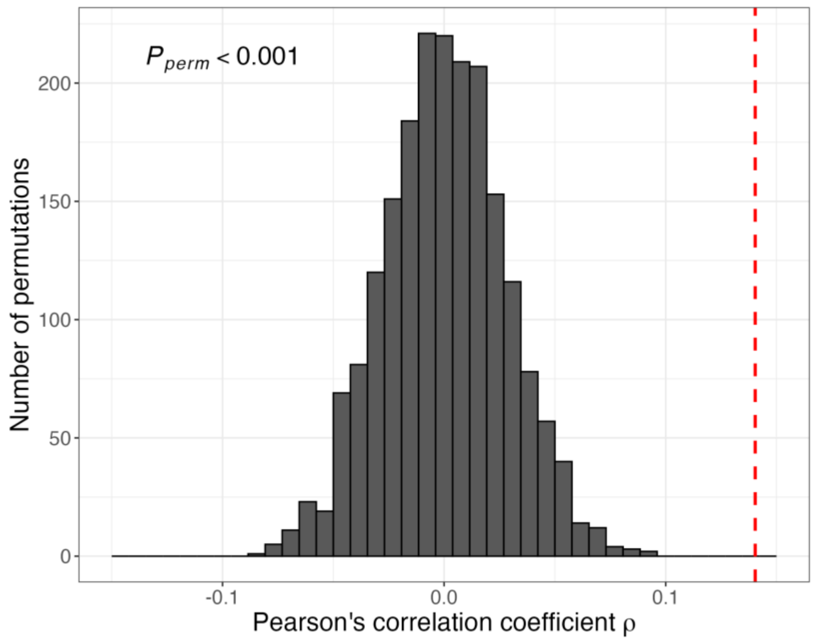


**Figure S3. Association between local densities of *M. reukaufii* and *A. nectaris* after removing uncountable local populations (n = 2,552).** Histogram shows the null distribution of Pearson correlation coefficients generated from 2,000 block-structured permutations to account for statistical non-independence (see Methods). The observed coefficient ($\rho = 0.151$, dashed vertical line) fell above the entire null distribution ($P < 0.001$).

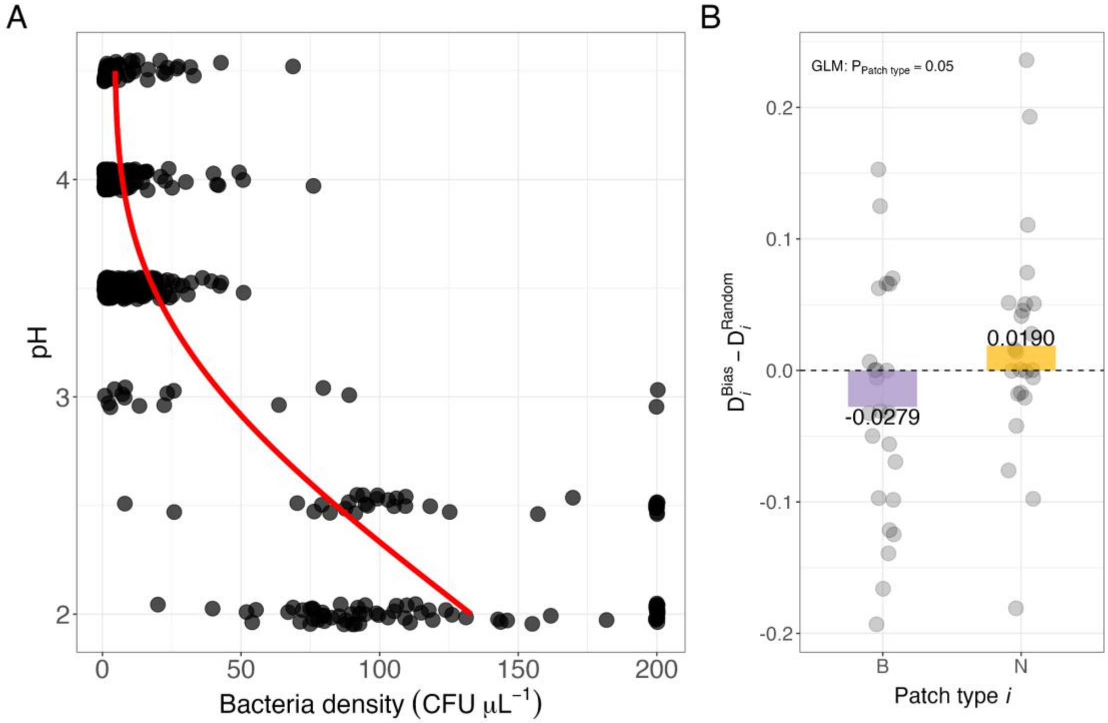


**Figure S4. Verifying presumed effects of experimental treatments.** (**A**) Addition of bacterial cells reduces the pH of nectar. The red curve was fitted via a generalised additive model (GAM; k = 3) purely to aid visualising the reduction of nectar pH by bacteria. (**B**) Dispersal propensities differ between bias and random dispersal treatments by patch type**.** Points indicate the difference in patch-specific dispersal propensity between a bias and random replicate, where a patch is either inoculated with bacteria (patch $B_+$) or not (patch $B_-$). See methods for details on how the metric was derived.

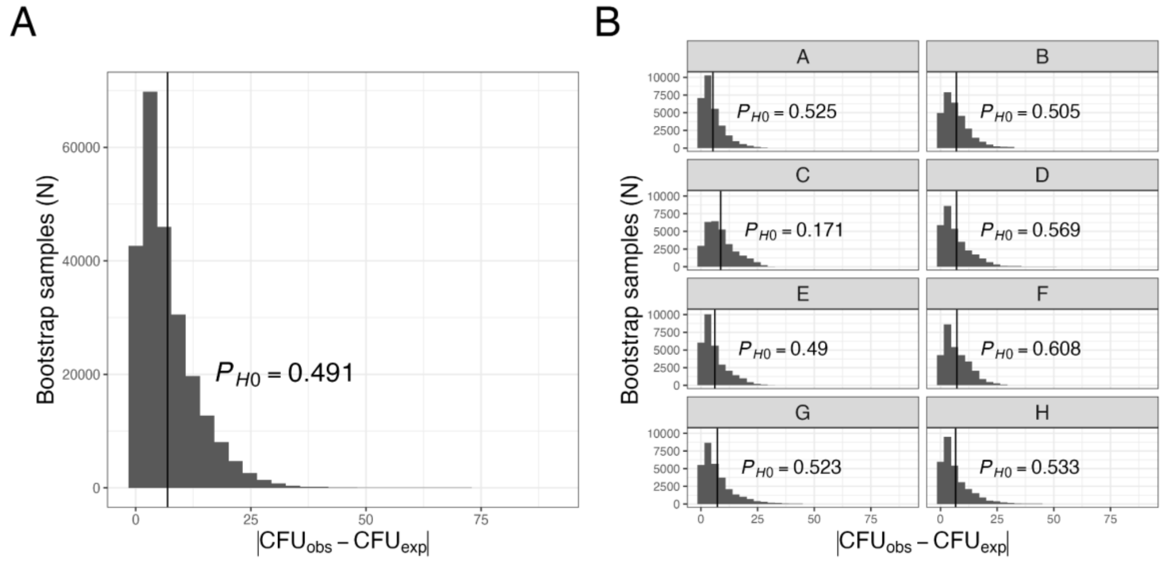


**Figure S5. Effect of microplate well position on yeast density.** Histograms show the stratified bootstrap distribution of the absolute difference between the observed average yeast density and the expected average within treatment/block. (**A)** shows the pooled deviations across all wells while (**B)** shows the well-specific deviances. The vertical line indicates the observed absolute difference between the yeast density of a well and its treatment-specific average. *P*-values represent the probability of observing a value at least as extreme as the observed value (vertical line) given that the null hypothesis is true. Since we did not correct for multiple testing in the well-specific tests (**B**), *P*-values are anti-conservative (*i.e.*, more likely to detect difference).

### Supplementary methods

#### Diplacus-like artificial nectar

Artificial nectar used in the experiment were made to resemble *D. aurantiacus* nectar in sugar and amino acid composition (Chappell et al., 2022; Peay et al., 2012). However, this protocol may have left some amino acids found in *D. aurantiacus* nectar missing in the artificial nectar used here (Vannette & Fukami, 2018), which may have caused *Acinetobacter nectaris* to behave as sink populations in our experimental system. Specifically, fructose (4%) glucose (2%), sucrose (20%), serine (0.102mM), glycine (0.097 mM), proline (0.038 mM), glutamate (0.035 mM), aspartic acid (0.026 mM), GABA1 (0.023 mM), and alanine (0.021 mM), were mixed until dissolved, and filtered through 0.2 µm filter. See **Artificial nectar protocol** for details.

#### Microbe strains and culturing

Our experiments used glycerol stocks of the bacterium *Acinetobacter nectaris* (strain FNA17) and the yeast *Metschnikowia reukaufii* (strain MR1) stored at -80°C. Both species were isolated from the floral nectar of *D. aurantiacus* in Jasper Ridge Biological Preserve, in 2017 (FNA17) and 2010 (MR1), respectively. *A. nectaris* was streaked on tryptic soy agar (TSA) containing 100 mg/L of cycloheximide and *M. reukaufii* on yeast mold agar (YMA) containing 100 mg/L of chloramphenicol. Streaked plates were then incubated at 26 °C. After 3-4 days of growth on agar plates, we resuspended cells of each species in their respective broth media (see **Broth culture**). After vortexing to homogenise, we aliquoted 50 µl of the suspended cell cultures by diluting them into several new 8-strip PCR tubes containing 50 µl of 50% glycerol. This gave us several 100 µl cultures of 25% glycerol stocks of each species, where each species was grown from the same initial culture to minimise genetic variability. These stock cultures were then stored at -80°C and used subsequently for the rest of the experiment.

#### Inoculum preparation

Inoculums for our main experiment were prepared by transferring 100 µl of glycerol stock cultures into 5 ml of broth media (YB and TSB for yeast and bacteria respectively) and culturing them for approximately 23 hours in a shaking incubator at 26°C and 120 RPM. To minimise bias introduced by inoculating our experimental plates with broth media, we washed inoculums by centrifuging them in 1mL eppendorf tubes (settings: 7.5min, 6k RPM), discarding the supernatant via pipetting, and resuspending cells in artificial nectar (i.e., experimental media) by vortexing. Lastly, the resulting overnight cells dissolved in nectar were then serially diluted by factors of $10^{-3}$ and $10^{-1}$, corresponding to approximately ~150 and ~10000-20000 cells/µl for *M. reaukaufii* and *A. nectaris* respectively.

#### Artificial nectar protocol

Ingredients table:

| Reagent | FW | Manufacturer | Product # | Lot # |
|---|---|---|---|---|
| (L) Serine | 105.09 | Sigma | S4311-25g | SLBT8375 |
| Glycine | 75.07 | Sigma | 50046-50g | SLBS3667 |
| (L) Proline | 115.1 | Sigma | P-0380 | 40F-0065 |
| (L) Alanine | 89.09 | Sigma | A7469-25g | SLB3605 |
| (L) Aspartic Acid | 133.10 | Sigma | A8949-25g | SLBP9902V |
| (L) Glutamine | 146.15 | Sigma | G8540-25g | 048K0074 |
| GABA1 (γ-Aminobutyric acid) | 103.12 | Sigma | A5836-10g | BCBS2563V |
| Sucrose | 342.2965 | Fisher Science Education | S25590B | 7GC0000150092 |
| Fructose | 180.15 | Fisher Science Education | S25332A | 6GCAF14031512 |
| Glucose | 180.15 | EMD Millipore | 346351-1KG | 2894283 |

**Protocol for making 200 mL:**

1. **Sugar solution**. Dissolve sugars in 200 ml DI water. Weights to dissolve and associated concentration:
   a. Fructose - 8.65 g (43.26 mg/ml)
   b. Glucose - 3.80 g (24.01 mg/ml)
   c. Sucrose - 39.70 g (198.49 mg/ml)
2. **Single amino acid stock solutions**. Make single 10ml solutions of each amino acid and refrigerate for later use. Weights to dissolve (in 10ml DI water) and associated concentrations:
   a. Serine (SER1) - 1.051 g (1 M)
   b. Glycine (GLY) - 0.751 g (1 M)
   c. Proline (PRO) - 1.151 g (1 M)
   d. Glutamine (GLU) - 0.059 g (0.04 M)
   e. Aspartic Acid (ASP) - 0.013 g (0.01 M)
   f. GABA1 - 1.031 g (1 M)
   g. Alanine (ALA) - 0.891 (1 M)
3. **Add amino acids to the sugar solution**. This is the nectar solution used for the experiment. Pipette amino acid solutions into the sugar solution to get the desired concentrations of AA in nectar. The volume to pipette into the 200 mL sugar solution and their final concentration in the nectar:
   a. Serine (SER1) - 20.4 µl (0.102 mmol)
   b. Glycine (GLY) - 19.4 µl (0.097 mmol)
   c. Proline (PRO) - 7.6 µl (0.038 mmol)
   d. Glutamine (GLU) - 175 µl (0.035 mmol)
   e. Aspartic Acid (ASP) - 520 µl (0.026 mmol)

f. GABA1 - 4.6 µl (0.023 mmol)
g. Alanine (ALA) - 4.2 µl (0.021 mmol)

4. **Sterilise nectar solution.** Sterilise the solution by filtering through 0.2 µm filter (Nalgene Rapid-Flow) inside a flow hood and refrigerate.

#### Agar plates

The recipe below is for 1000 ml solutions. Dissolve ingredients in 1000 ml DI water and autoclave before plating onto approximately 20 plates (100mm x 15mm petri dish).

**Yeast mold agar 1L (YMA)**

1. Malt extract - 3 g
2. Bactopeptone - 5 g
3. D(+) glucose (deoxytrose) - 10 g
4. Bacteriological agar - 20g
5. Yeast extract - 3g

**Tryptic soy agar 1L (TSA)**

1. Tryptic soy agar - 40 g

#### Broth culture

Dissolve ingredients in 100 ml of DI water and autoclave.

**Yeast broth**

1. Malt extract - 0.3 g
2. Bactopeptone - 0.5 g
3. D(+) glucose (deoxytrose) - 1 g
4. Yeast extract - 0.3 g

**Bacteria broth**

1. Tryptic soy broth - 3g

**Sample image of colony forming units on yeast (top) and bacteria (bottom) agar plates**

- Images show plate 5, dilution cycle 4 (23Apr23)
- Each row of CFUs corresponds to a single treatment
- Bacteria cells tend to stay in the wells that they were added into (A-C or A-F)

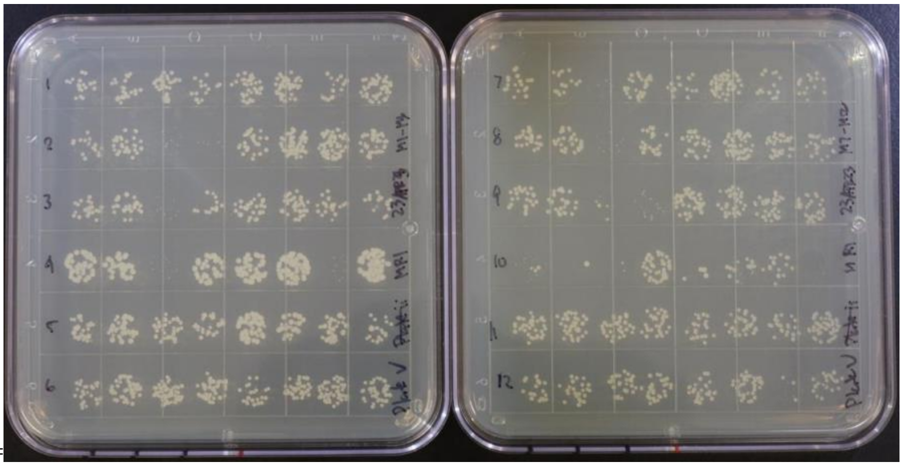

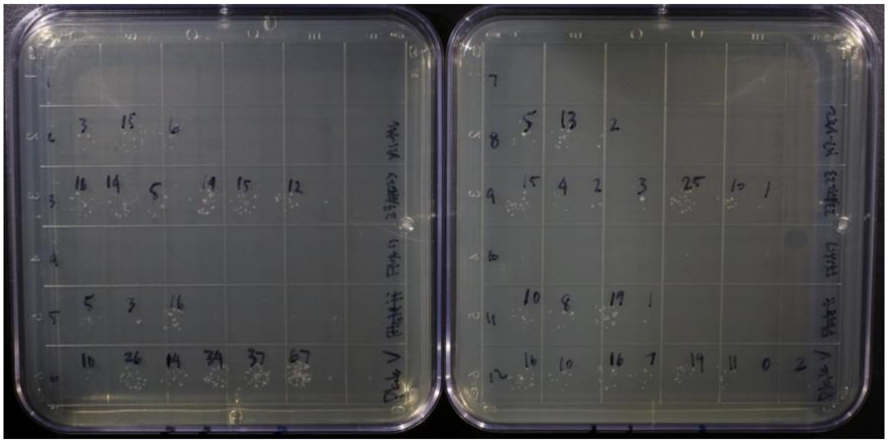

**Statistical tests of experimental artefacts**

*Type-specific dispersal propensity*

To confirm that our dispersal bias treatment reduced the dispersal rate of bacteria inoculated patches (and increased those without bacteria inoculation), we derived a metric of dispersal propensity from the dispersal sequence data generated from our experiment. Because the number of dispersal events within a single time point is sparse (3 or 6), we computed dispersal propensities by aggregating the data across the duration of the experiments. Specifically, the complete dispersal sequence data of a replicate metapopulation is stored as an edge list, specifying the patches that nectar was transferred from (i.e., emigration/donor) and to (i.e., immigration/recipient). For each metapopulation edge list, we tabulated the number of times each patch type (i.e., bacteria added (B), no bacteria added (N)) was selected for emigration and immigration. The total emigration and immigration events per type were then normalised by the total dispersal events (12 or 24) to make replicates with different dispersal frequencies comparable. We then took the product of these normalised emigration and immigration rates as a measure of dispersal propensity. Because what mattered is that the dispersal propensities differed between the bias and random dispersal treatments (i.e., pollinators prefer patches with less acidic nectar), we took the difference between dispersal propensities of the bias and random dispersal treatments, denoted as $D^{bias}$ and $D^{random}$, respectively. Thus, if the dispersal bias treatment worked, the difference in dispersal propensities should be <0 and >0 for bacteria ($D_B^{bias}$ - $D_B^{random}$) and no bacteria patches ($D_N^{bias}$ - $D_N^{random}$), respectively. As a reminder, dispersal is defined at the patch level (and not microbe specific) because pollinators indirectly move microbes by transferring nectar between flowers.

*Effect of microplate well position*

To minimise experimental errors, we added bacterial sink populations to the same well across all replicates. To ensure that our results are not artefacts of the choice of wells inoculated with bacteria, we tested the effects of well position on yeast densities through a stratified bootstrapping procedure that partials out the effects of all experimental treatments (e.g., dispersal frequency, dispersal bias, sink addition, and date). Specifically, within each treatment block, we first calculated the mean yeast density across replicates for each well position (A through H), which served as the observed value (*i.e.*, $CFU_{obs}$). We then constructed a null distribution assuming equal yeast densities across wells by resampling yeast densities with replacement from the pooled distribution of all wells within the block and computing the mean for each resample (*i.e.*, $CFU_{exp}$). This procedure was repeated 500 times per well in each block. We then pooled all resampled values across blocks and wells (60 blocks × 8 wells × 500 samples = 240,000) to form the null distribution. The p-value was calculated as the proportion of null-generated mean densities that were at least as extreme as the observed mean density.

**Work cited**

Vannette, R. L., & Fukami, T. (2018). Contrasting effects of yeasts and bacteria on floral nectar traits. *Annals of Botany*, *121*(7), 1343–1349. https://doi.org/10.1093/aob/mcy032
